# Ingestion of *Bacillus cereus* spores dampens the immune response to favor bacterial persistence

**DOI:** 10.1101/2023.03.16.532769

**Authors:** Salma Hachfi, Alexandra Brun-Barale, Patrick Munro, Marie-Paule Nawrot-Esposito, Gregory Michel, Arnaud Fichant, Mathilde Bonis, Raymond Ruimy, Laurent Boyer, Armel Gallet

**Affiliations:** Université Côte d’Azur, CNRS, INRAE, ISA, France; Université Côte d’Azur, Inserm, C3M, Nice, France; Anses (Laboratoire de Sécurité des Aliments), Université Paris-Est, France; Bacteriology Laboratory, Archet 2 Hospital, CHU, Université Côte d’Azur, Nice, France

**Keywords:** Intestine, *Bacillus* spores, Innate immunity, Amidases, *Drosophila*, Mice

## Abstract

Spores are considered as dormant entities highly resistant to extreme conditions. Among them, *Bacillus cereus* spores are commonly associated with foodborne outbreaks. Nevertheless, the pathological processes associated with spore ingestion and germination remain poorly understood. Here, we show that while ingestion of vegetative bacteria leads to their elimination from the midgut and small intestines of *Drosophila* and mice, respectively, a single ingestion of spores leads to the persistence of bacteria for at least 10 days. Using *Drosophila* genetics, we demonstrate that spores escape the innate immune response of the anterior midgut. Once in the posterior midgut, spores germinate, and the vegetative cells dampen the immune signaling through the induction of amidases which are negative regulators of the immune response. This study provides evidence for how *B. cereus* spores hijack the intestinal immune defenses allowing the localized birth of vegetative bacteria responsible for the digestive symptoms associated with foodborne illness outbreaks.

## INTRODUCTION

Organisms are subjected to various environmental stresses including starvation, temperature variation, chemicals and microbes. Healthy individuals overcome these assaults by engaging defense mechanisms that maintain their homeostasis. Among the stressors, opportunistic enteric bacteria become pathogenic when host defenses are diminished or inefficient.

The evolutionarily conserved innate immune system is the first line of defense against bacteria, and adult *Drosophila* has proven to be a powerful model for innate immunity studies (Capo et al., 2019). In the midgut, the local innate immune system is mainly mounted in the anterior parts (Capo et al., 2019; Royet and Charroux, 2013). First, anterior enterocytes can rapidly sense the presence of allochthonous (i.e., non-commensal) bacteria and secrete reactive oxygen species (ROS) in a DUOX-dependent manner to block bacterial proliferation (Benguettat et al., 2018; Kim and Lee, 2014). Concomitantly, luminal ROS are perceived by a subpopulation of anterior enteroendocrine cells that respond by releasing the DH31/CGRP neuropeptide, which promotes visceral muscle spasms to provoke the expulsion of bacteria from the midgut in less than 4 hours post-ingestion (Benguettat et al., 2018). Nevertheless, if the load of allochthonous bacteria is higher and/or if the immune ROS and visceral spasms are insufficient to eliminate them, the allochthonous bacteria can start to proliferate, releasing muropeptides from the peptidoglycan (PGN), a bacterial cell wall component, that bind to the transmembrane and intracellular immune receptors PGRP-LC and PGRP-LE, respectively (Bosco-Drayon et al., 2012; Kaneko et al., 2006). Consequently, the Immune deficiency (Imd)/NF-κB pathway is activated, leading to the expression of anti-microbial peptides (AMPs) beginning 4-6 hours post-ingestion (Bonfini et al., 2016; Buchon et al., 2009b; Chakrabarti et al., 2012; Zhai et al., 2018b). Because a prolonged activation of the local innate immunity is responsible for chronical inflammation which is detrimental for the individual, a robust negative feedback is turned on to tune down the Imd pathway once the bacteria are cleared (Zhai et al., 2018b). For instance, the PGRP-SC1 & 2 and PGRP-LB amidases have been described as negative regulators of the Imd pathway mainly acting by digesting the extracellular PGN fragments and thus blocking the recognition process by the Imd pathway receptors (Costechareyre et al., 2016; Guo et al., 2014; Kaneko et al., 2006; Paredes et al., 2011). Altogether, these combined means of defense allow an efficient sensing and elimination of the allochthonous bacteria allowing the survival of the individual.

However, the detection of bacterial spores by the local innate immune system of the intestine and their elimination remains challenging for host organisms. Indeed, spores can resist many biological, chemical, or physical treatments (Ehling-Schulz et al., 2019; Setlow, 2014). Among opportunistic bacteria, the widespread environmental spore-forming bacteria belonging to the *Bacillus cereus* (*Bc*) group are well-known worldwide food poisoning pathogens that cause diarrheal and/or emetic-type illnesses (Dietrich et al., 2021; Jovanovic et al., 2021). *Bc* is the second most important cause of foodborne outbreaks (FBOs) in France (Bonis et al., 2021; Glasset et al., 2016; Santé publique France, 2019) and the third in Europe (EFSA and ECDC, 2018). When nutrient-rich conditions are encountered, *Bc* spores can germinate, giving rise to vegetative cells which can even proliferate. Such favorable conditions are present in the small intestine of mammals, where it is assumed that spores are able to germinate and probably proliferate to ultimately trigger diarrhea due to the production of enterotoxins (Berthold-Pluta et al., 2015; Ehling-Schulz et al., 2019). *In vitro* experiments have indicated that *Bc* vegetative cells can be destroyed by the acidic pH of the stomach and the biliary salt of the duodenum while spores can resist (Barbosa et al., 2005; Berthold-Pluta et al., 2015; Ceuppens et al., 2012a; Ceuppens et al., 2012b; Ceuppens et al., 2012c; Clavel et al., 2004). The effectiveness of the gut innate immune system to fight *Bc* vegetative cells has been demonstrated (Benguettat et al., 2018). In contrast, nothing is known concerning the behavior and the fate of *Bc* spores in the intestine *in vivo* and the related immune response mounted by the host.

Here, thanks to *in vivo* studies, we demonstrate that spores germinate only in the posterior parts of the *Drosophila* midgut and of the mouse small intestine. We show that bacteria of *Bc* group can persist for at least ten days in the posterior regions of the midgut/small intestine. Next, we demonstrate that the spores do not trigger any detectable immune responses in the anterior parts of the *Drosophila* midgut. Once in the posterior midgut, germinated cells trigger the Imd immune pathway in a PGRP-LE dependent manner. Strikingly, we found the amidases PGRP-SC1/2 and PGRP-LB, which are negative regulators of the Imd pathways, to be induced while the AMPs were repressed. In flies deficient in the PGRP-LE receptor, the cytosolic members of the Imd pathway or the PGRP-SC1/2 and PGRP-LB amidases, the persistence of the bacteria in midgut was reduced. Surprisingly, removing Relish, the NF-κB-like transcription factor, downstream of the Imd pathway has no impact on the bacterial persistence. However, depletion of Dif, another NF-kB-like transcription factor, together with Relish provided the proof of a critical cooperation of both transcription factors in regulating *amidase* and *AMP* expressions in the posterior midgut and thus the bacterial clearance. Altogether, we provide evidence that spores belonging to the *Bc* group persist in the intestine when ingested as spores and escape the anterior immune response. Spores reach the posterior regions of the midgut where they germinate, and vegetative bacteria induce expression of amidases that act as negative regulators of the Imd pathway, dampening the production of AMPs, consequently fostering the persistence of the bacteria.

## RESULTS

### Spores of the *Bc* group persist in the *Drosophila* intestine

The *Bc* group is subdivided into at least eight phylogenetically very close genomospecies (Carroll et al., 2021; Carroll et al., 2020; Ehling-Schulz et al., 2019). For this study, we selected the reference strain *Bc sensu stricto (Bc* ATCC 14579) as well as two *Bacillus thuringiensis* subspecies *kurstaki* (*Btk*) strains (*Btk* SA-11 and *Btk* ABTS-351) because of their broad use as microbial pesticides and the fact that *Btk* has been suspected to be responsible for FBOs (Biggel et al., 2021; Bonis et al., 2021; Johler et al., 2018).

To determine whether, upon ingestion, *Bc* or *Btk* spores behaved the same as vegetative cells, we fed flies continuously with contaminated food and assessed the amount of *Bc* or *Btk* bacteria in the *Drosophila* midgut at different times (Figure 1A). We observed that regardless of the strain used, *Bc*/*Btk* were still present in the *Drosophila* midgut at least 10 days after the initial contact with spore-contaminated food (Figure 1B).

**Figure 1.**
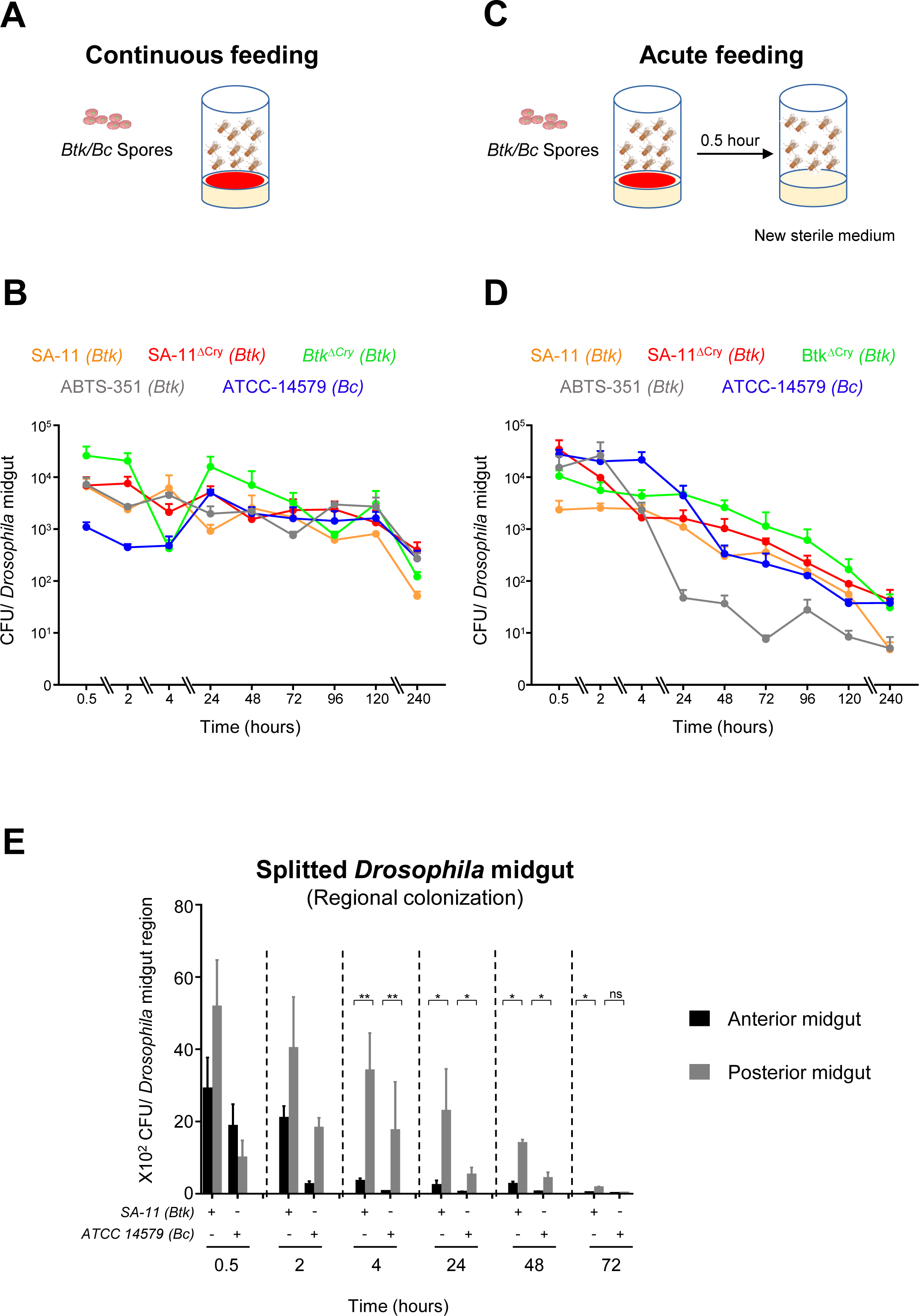
Spores of the *Bacillus cereus* group persist in the *Drosophila* intestine. **(A)** Experimental setup to assess bacterial load after a continuous ingestion of spores. **(B)** Bacterial loads of dissected midguts after continuous ingestion of spores from *Btk* or *Bc* strains. The vertical axis indicates the median number of colony-forming units (CFUs) per midgut of at least three independent experiments. Error bars correspond to the SEM. **(C)** Experimental setup to assess bacterial load after an acute ingestion of spores. Flies are in contact with the contaminated medium for 30 minutes and then transferred to fresh vial devoid of spores. **(D)** Bacterial loads of dissected midguts after acute ingestion of spores from *Btk* or *Bc* strains. The vertical axis indicates the median number of CFUs per midgut of at least four independent experiments. Error bars correspond to the SEM. **(E)** Bacterial loads in split *Drosophila* midguts after acute intoxication with *Btk* (SA-11) or *Bc* spores. Error bars correspond to the SEM of at least three independent experiments. Mann-Whitney tests were applied. Asterisks represent a statistically significant differences between bacterial loads in the anterior and the posterior midguts: **p < 0.01, *p < 0.05

These data were very different from what we observed using *Bc* ATCC 14579 or *Btk* SA-11 vegetative cells, whose loads in the intestine rapidly decreased within 24 hours (Figure S1A). Unlike *Bc*, during sporulation, *Btk* produces Cry toxins embedded in a crystal, which displays specific entomopathogenic properties. *Btk* is widely used specifically to kill lepidopteran larvae that are broad crop pests (Ehling-Schulz et al., 2019). Because the presence of Cry toxins might influence the behavior of *Btk* in the midgut, we engineered a *Btk* SA-11*^ΔCry^* strain cured of its plasmids and therefore devoid of Cry toxins (see Experimental Procedures). We also used a *Btk ^ΔCry^* obtained from the Bacillus Genetic Stock Center (#4D22, https://bgsc.org/) also devoid of Cry toxins. Importantly, the SA-11*^ΔCry^* and the *Btk^ΔCry^* spores behaved like *Btk* SA-11 and *Bc* ATCC 14579 spores (Figure 1B), refuting any possible role of the Cry toxins in this phenomenon. The commercially available preparation of the *Btk* ABTS-351 spores is known to contain 46% of additives (ec.europa.eu). To assess the potential involvement of those additives in the intestinal persistence of *Btk* ABTS-351, we extended our study to the commercial preparation, which we compared with a *Btk* ABTS-351 spore preparation made in our laboratory without any additives. We noticed that the commercial spores as well as the “homemade” *Btk* ABTS-351 spores behave similarly in the *Drosophila* midgut (Figure 1B and S1B), suggesting that the additives present in the commercial preparation did not contribute to the persistence of *Btk* ABTS-351 spores.

Because spores could germinate and proliferate on the fly medium, we checked this possibility by counting the number of *Btk* SA-11 bacteria on the fly medium in the absence of flies. We applied a heat treatment in order to kill all germinated vegetative cells. We noticed that 2 days after spore deposit on the fly medium, some of them started their germination and even proliferated 4 days after deposit (Figure S1C). Hence, to remove this limitation in the persistence assessment, we performed acute feeding. Flies were fed with spores for 30 min before being transferred onto fresh food medium (i.e. without spores) (Figure 1C). We first verified that upon acute ingestion of *Bc* ATCC 14579 or *Btk* SA-11 vegetative cells, they were readily cleared from *Drosophila* midguts (Benguettat et al., 2018) (Figure S1D). We then monitored the persistence of spores in the *Drosophila* midgut upon acute feeding (Figure 1D). After 30 min of spore feeding, the bacterial load averaged 10^4^ cells per midgut regardless of *Bc*/*Btk* strain. Interestingly, the bacteria could persist up to 10 days in the *Drosophila* midgut after acute spore ingestion (Figure 1D and S1E). The same observation was made with *Btk* SA-11 spores in a different *Drosophila* genetic background (i.e., *w^1118^ Drosophila* midgut, Figure S1F). These results demonstrate that, unlike ingestion of vegetative cells, ingestion of *Bc*/*Btk* spores results in bacterial persistence for several days in the *Drosophila* midgut.

The *Drosophila* midgut is subdivided into five major anatomical regions (R1 to R5) (Figure S1G) (Buchon et al., 2013; Marianes and Spradling, 2013). To analyze in detail the localization of *Bc*/*Btk* along the midgut, we quantified the bacterial load in the anterior and posterior midgut after acute feeding. We did not focus on the acidic region due to its small size and the difficulty to dissect it accurately. During the first two hours after acute ingestion, we found that *Bc*/*Btk* bacteria were present in both regions of the midgut (anterior and posterior). However, from 4 hours onward, the posterior midgut harbored a significantly higher load of *Bc*/*Btk* cells compared with the anterior midgut (Figure 1E). Collectively, our results demonstrate that *Bc*/*Btk* persist for up to 10 days in the midgut and may accumulate preferentially in the posterior regions.

### Spores of the *Bc* group germinate preferentially in the posterior midgut

Spores are metabolically dormant and resistant to extreme environmental conditions, allowing them to survive to extreme conditions (Setlow, 2014). However, the presence of nutrients can trigger the process of germination, in which spores emerge from dormancy, growing into vegetative cells. Since the intestine is a favorable environment for spore germination, we hypothesized that *Bc*/*Btk* spores might germinate in the *Drosophila* midgut. To address this question, we developed a robust fluorescent staining technique suitable for visualizing spores and differentiating them from their outgrowing vegetative cells. Dormant *Bc*/*Btk* spores harbored a red fluorescence. Once germinated, vegetative cells started to express the green fluorescent protein (GFP). The use of this novel tool endowed with dual red/green (R/G) labelling allowed us to follow the process of germination in real time (Figure 2A, movie1).

**Figure 2:**
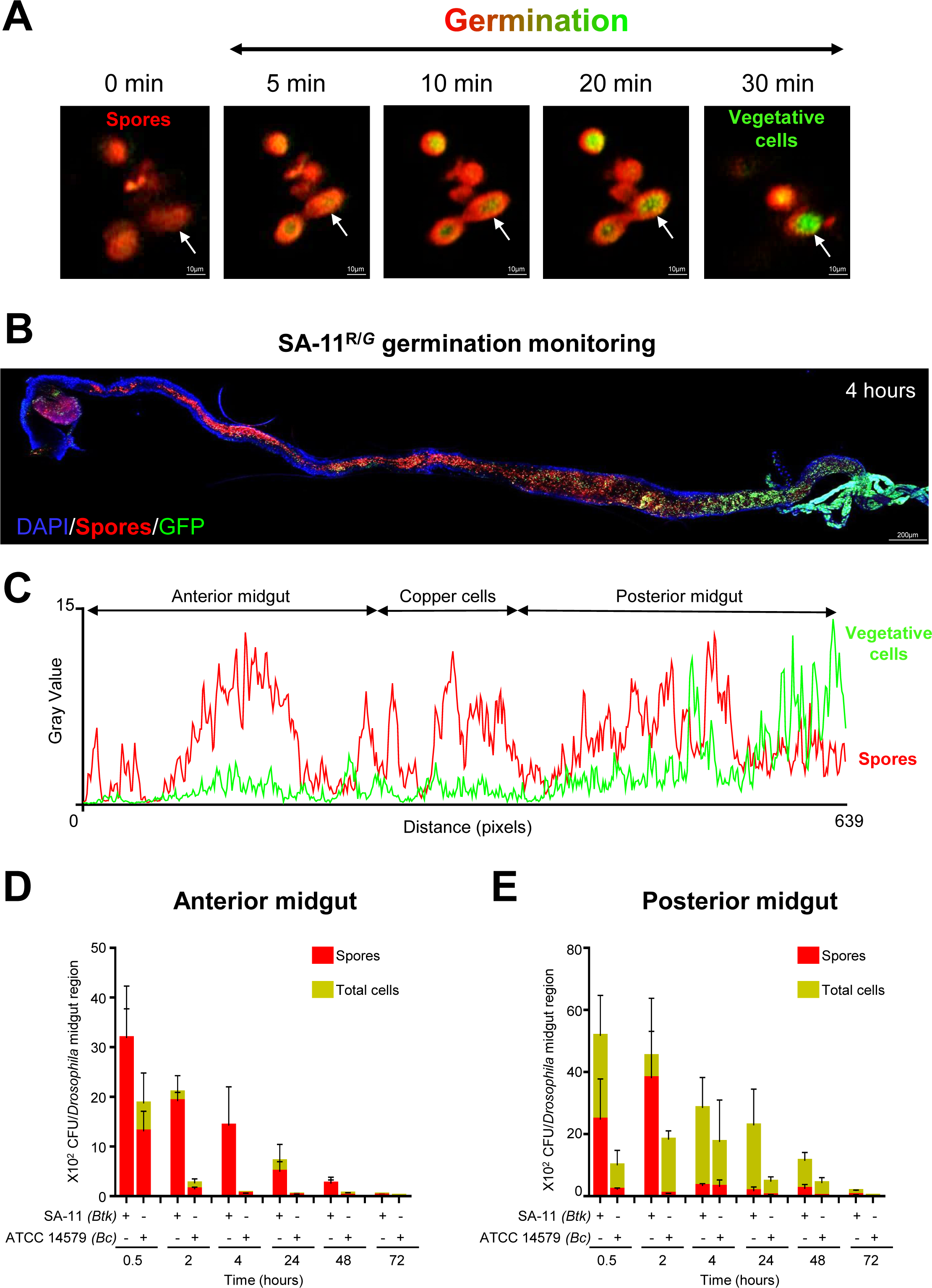
SA-11 spores germinate preferentially in *Drosophila* posterior midgut. **(A)** Time-lapse images of SA-11*^R/G^* spores during germination. **(B)** Monitoring of SA-11*^R/G^* germination *in vivo* in *Drosophila* midgut 4 h post-spore-ingestion. **(C)** Plots of the average fluorescence intensity (represented as mean gray value) of SA-11^R/G^ germination measured along the *Drosophila* midgut presented in (B). For all plot analyses of average fluorescence shown in this paper, the red line represents the average of the spore fluorescence (Red) and the green line represents the average fluorescence of vegetative cells (GFP). **(D-E)** Bacterial load of SA-11 or *Bc* in anterior (D) and posterior (E) *Drosophila* midguts. Green bars (non-heated samples) represent the whole SA-11 or *Bc* bacterial loads (spores and vegetative cells). Red bars (heated samples) represent the proportion of spore loads. Data represent the mean ± SEM of at least five independent experiments.

The use of the *Btk* SA-11*^R/G^* fluorescent strain *in vivo* first revealed that at 0.5− and 2-hours post-ingestion, *Btk* spores occupied the lumen of the whole midgut (Figure S2A and S2B). Few vegetative cells were detectable in the posterior midgut 2 hours post-ingestion (inset in Figure S2B). *Btk* SA-11*^R/G^* spore germination was evident in the posterior midgut 4 hours after ingestion (Figures 2B and 2C). Interestingly, we found that regardless of the *Bc* group strain, spore germination occurred markedly in the *Drosophila* posterior midgut at 4 hours after ingestion (Figure S2D). Twenty-four hours post-ingestion, we detected mostly vegetative cells in the posterior midgut (Figure S2C). To further confirm that spores mainly germinated in the posterior midgut, we performed measurements of colony-forming units (CFUs) in the anterior and posterior parts of the midgut by comparing heat-treated intestinal samples (to kill germinating spores and vegetative cells but not spores) to non-heat-treated samples (cumulating spores, germinating spores and vegetative cells). Interestingly, we did observe in the anterior midgut region the almost exclusive presence of *Bc*/*Btk* spores, even 3 days after acute ingestion (Figure 2D). However, in the posterior midgut, we observed the appearance of the first *Bc*/*Btk* germinating spores as early as 30 minutes after ingestion (Figure 2E). Together, these results demonstrate that the germination of *Bc* group spores begins 30 minutes after oral ingestion and occurs mainly in the posterior midgut of *Drosophila melanogaster*.

### Spores do not trigger detectable *Drosophila* midgut innate immune response

The persistence of spores in the *Drosophila* midgut raises the question of how the local innate immune system can tolerate spores and/or vegetative cells. As mentioned previously, in response to enteric infection, the anterior *Drosophila* midgut (R1 and R2 regions) initiates immune responses via the luminal release of ROS and, if necessary, AMPs (Benguettat et al., 2018; Bosco-Drayon et al., 2012; Hachfi et al., 2019; Lee et al., 2013; Royet and Charroux, 2013; Tzou et al., 2000). To test the release of local immune ROS (HOCl) in response to *Btk* vegetative cells or spores, we used the R19S probe, a HOCl sensitive fluorescent dye (Chen et al., 2011; Hachfi et al., 2019). First, we confirmed that *Btk* vegetative cells induced ROS only in the anterior region one-hour post-ingestion (Figure 3A, middle panel). Surprisingly, *Drosophila* fed with *Btk* spores did not show ROS induction either in the anterior or in the posterior regions of the *Drosophila* midgut (Figure 3A, lower panel), though spores germinated in the posterior midgut (Figures 2 and S2). In parallel, we specifically knocked down the expression of *Duox* in *Drosophila* enterocytes by RNA interference and examined the resulting impact on the spore persistence. We first confirmed that, 4 hours after ingestion of vegetative cells, the silencing of *Duox* in the enterocytes increased the load of *Btk* compared with control intestines (Benguettat et al., 2018) (Figure 3B). However, *Duox* silencing in *Drosophila* enterocytes did not impact the *Btk* persistence after spore ingestion (Figure 3B). Based on these data, we inferred that ingestion of spores does not induce the production of *Duox-*dependent ROS.

**Figure 3:**
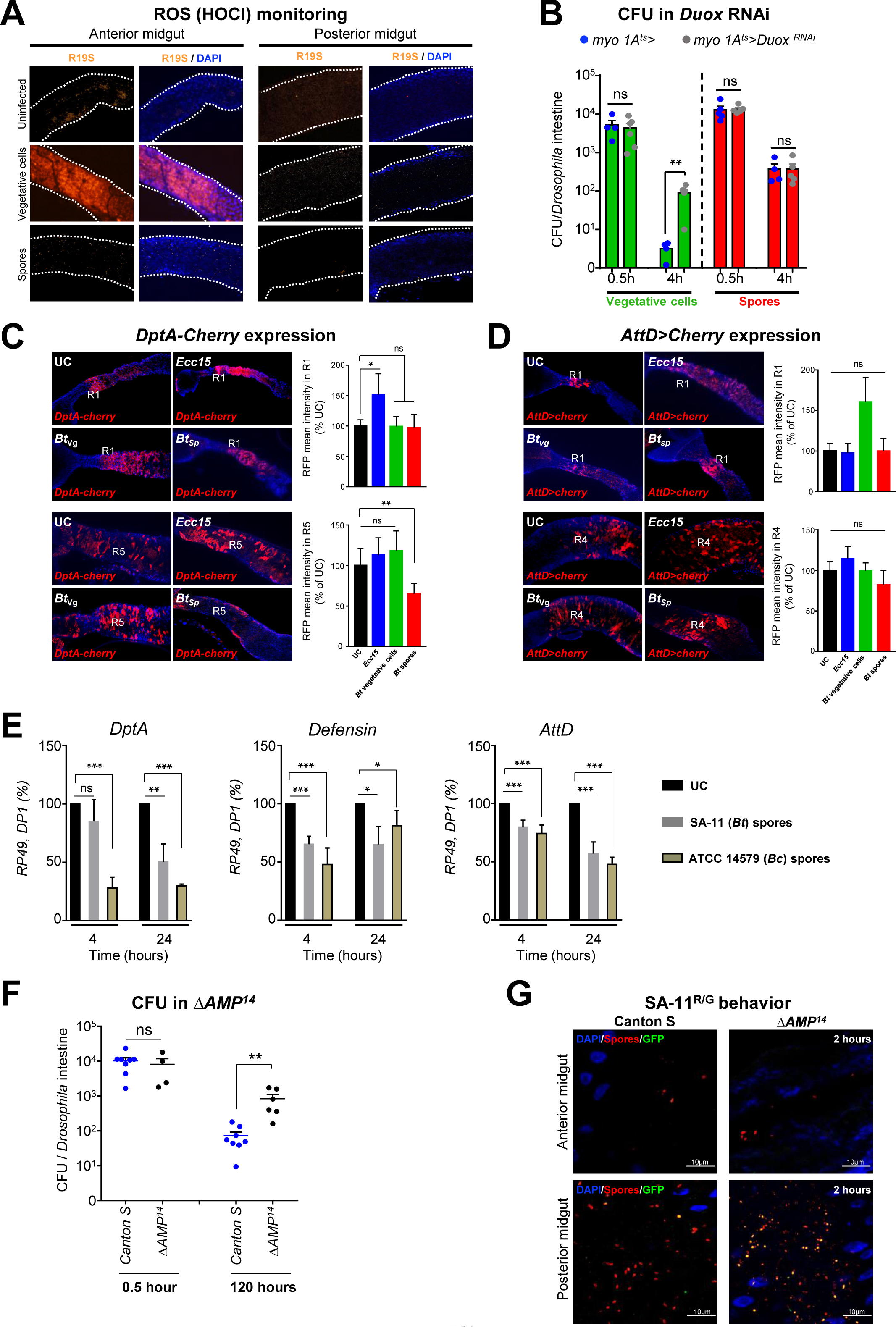
*Bacillus cereus* spores do not trigger midgut innate immune response. **(A)** ROS monitoring after acute feeding with SA-11 spores or vegetative cells. ROS production in the midgut is visualized by the HOCl-specific R19S probe (orange). DAPI (blue) marks the nuclei. **(B)** SA-11 loads in midguts knocked down for the expression of *Duox* in enterocytes 0.5 or 4 hours after acute feeding with vegetative cells or spores. The horizontal axis indicates the median number of CFUs per midgut of at least three independent experiments. Error bars correspond to the SEM. **(C)** *DptA-Cherry* expression (red) in the R1 midgut region (upper panel) and in the R5 midgut region (bottom panel) of *Drosophila* fed for 30 minutes with H_2_O, *Ecc15*, SA-11 vegetative cells (*Bt_vg_*) or SA-11 spores (*Bt_sp_*) and observed 24 hours later. Measured quantities are shown on the right graphs. The results are given as the relative expression compared with the control (H_2_O). Data represent means ± SEM of at least three independent experiments. **(D)** *AttD-Gal4 UAS-Cherry* expression (red) in the R1 midgut region (upper panel) and in R4 midgut region (bottom panel) of *Drosophila* fed for 30 minutes with H_2_O, *Ecc15*, SA-11 vegetative cells (*Bt_vg_*), or SA-11 spores (*Bt_sp_*) and observed 24 hours later. Measured quantities are shown on the right graphs. The results are given as the relative expression compared with the control (H_2_O). Data represent means ± SEM of at least three independent experiments. **(E)** qRT-PCR analyses of *AMP* expression in midgut upon acute feeding with SA-11 or *Bc* spores. UC corresponds to flies fed with water. For RT-qPCR results, mRNA levels in unchallenged wild-type flies were set to 100 and all other values were expressed as a percentage of this value. RT-qPCR results are shown as mean ± SEM from 10 female flies per genotype from at least three independent experiments. **(F)** Bacterial load in the midguts of Δ*AMP^14^* mutant flies 0.5 or 4 hours after acute feeding with SA-11 spores. The vertical axis indicates the median number of CFUs per midgut. Error bars correspond to the SEM. **(G)** Representative confocal images showing SA-11^R/G^ spore germination in the anterior and posterior midgut of WT (Canton S) and Δ*AMP^14^* mutant flies 6 hours after acute feeding with spores. DAPI (blue) marks the nuclei. Spores are in red, vegetative cells in green. The yellow fluorescence corresponds to germinating spores (see Figure 2A). The Mann-Whitney test was applied in B, C, D and F. Student’s t-tests were used to analyze data in E. *p≤0.05, **p≤ 0.01, ***p≤0.001, ns = non-significant.

We next investigated the induction of *AMP* genes in the *Drosophila* midgut following acute ingestion of vegetative cells vs. spores. Using the *DiptericinA-Cherry* (*DptA-Cherry*) and *AttacinD-Gal4 UAS-Cherry* (*AttD>Cherry*) reporters, two readouts for the activation of the Imd pathway in the midgut (Charroux et al., 2018), we first observed that the acute ingestion of vegetative cells of the *Erwinia carotovora carotovora* (*Ecc15*) opportunistic bacteria induced *DptA* and *AttD* expression in the anterior midgut (Figure 3C, 3D, S3A and S3B). *Ecc15* was also capable of promoting the spreading of *AttD* expression in the posterior R4 region (Figure S3B). However, *Btk* vegetative cells did not show significant changes in *DptA* expression in either the anterior or posterior midgut (Figure 3C and S3A), while *AttD* was slightly induced, though not significantly, in the anterior midgut (Figure 3D). These data are consistent with the fact that early ROS induction followed by the visceral spasms are sufficient to rapidly eliminate *Btk* vegetative cells upon acute ingestion (Figure 3A and 3B) (Benguettat et al., 2018), at least before the Imd pathway can be induced. RT-qPCR analyses of the expression of *DptA, Defensin and AttD* genes on dissected midguts confirmed the non-induction of *AMP* genes after acute ingestion of *Btk* (or *Bc*) vegetative cells (Figure S3C).

Monitoring *AMP* expression upon acute spore feeding revealed that neither *DptA-Cherry* nor *AttD-Cherry* reporter genes were induced *in vivo* in the anterior midgut (Figure 3C, 3D, S3A and S3B). Strikingly, repression of the *DptA-Cherry* reporter expression in the posterior midgut was observed (Figure 3C and S3A). RT-qPCR analyses confirmed the repression of *DptA* expression as well as the repression of *Defensin* and *AttD* expression (Figure 3E). Together these data suggest that *Bc/Btk* persistence upon spore ingestion could be supported by decreased expression of *AMPs*.

To assess the involvement of the AMPs in *Btk* SA-11 persistence, we used a fly strain in which the 14 *AMP* genes were deleted (Δ*AMP^14^*) (Carboni et al., 2021; Hanson et al., 2019). We quantified the *Btk* SA-11 load in the midgut of wild type (*Canton S*) and Δ*AMP^14^* flies. While both genotypes ingested a similar amount of spores during the 30 min of feeding (Figure 3F), 120h post-feeding, in Δ*AMP^14^* mutant flies, we found a significantly higher *Btk* SA-11 load compared with wild-type flies (Figure 3F). This result suggests that AMPs could kill germinating cells in the posterior midgut. To further challenge this hypothesis, we monitored the fate of the *Btk* SA-11*^R/G^* fluorescent strain in Δ*AMP^14^* mutant flies after acute ingestion of spores. As expected, two hours post-ingestion, confocal imaging showed higher levels of germinating spores in Δ*AMP^14^* mutant posterior midguts than in WT posterior midguts (Figure 3G). We did not observe obvious changes in the anterior midguts where only spores were present (Figure 3G). Collectively, our data suggest that, though repressed upon spore ingestion, the weak production of AMPs is necessary to limit the *Bc/Btk* bacterial load in the posterior midgut.

### Amidases contribute to the intestinal persistence of spores

Since *AMPs* expression was downregulated after spore ingestion, we wondered whether the amidases, which exert negative feedback on the Imd pathway, could be induced by germinating spores (Charroux et al., 2018; Costechareyre et al., 2016; Paredes et al., 2011). First, we verified by RT-qPCR analyses that the ingestion of *Bc/Btk* vegetative cells could induce the expression of the three *amidase* encoding genes in the midgut (Figure S4). Interestingly, after spore ingestion, we observed that PGRP-SC1 and −SC2 were consistently induced both 4 hours and 24 hours post-feeding while PGRP-LB was only induced at 4 hours and was repressed at 24h hours (Figure 4A). We next assessed the *Btk* SA-11 intestinal load in mutant flies homozygous for either the *PGRP-SC1/2* or *PGRP-LB* loss-of-function alleles (*PGRP-SC1/2^Δ^* and *PGRP-LB^Δ^* respectively) (Paredes et al., 2011). In midguts from these mutant animals, no difference in bacterial load was observed compared with control flies 30 min after spore feeding (Figure 4B). However, 120 hours post-feeding, the loss of *PGRP-SC1/2* or *PGRP-LB* was associated with a significant decrease in the number of *Btk* SA-11 in the midgut compared with the *Canton S* flies (Figure 4B). Because spores accumulated and germinated in the posterior midgut of *Canton S* flies as early as 4 hours post-ingestion (Figure 2), we monitored the fate of the spores in *PGRP-SC1/2^Δ^* and *PGRP-LB^Δ^* deficient flies. Six hours after acute ingestion, confocal imaging showed the presence of fewer red/green germinating spores in the posterior midguts of *PGRP-SC1/2*^Δ^ or *PGRP-LB^Δ^* flies compared with WT flies (Figure 4C), suggesting that in the absence of amidases, the production of AMPs was able to kill germinating spores.

**Figure 4:**
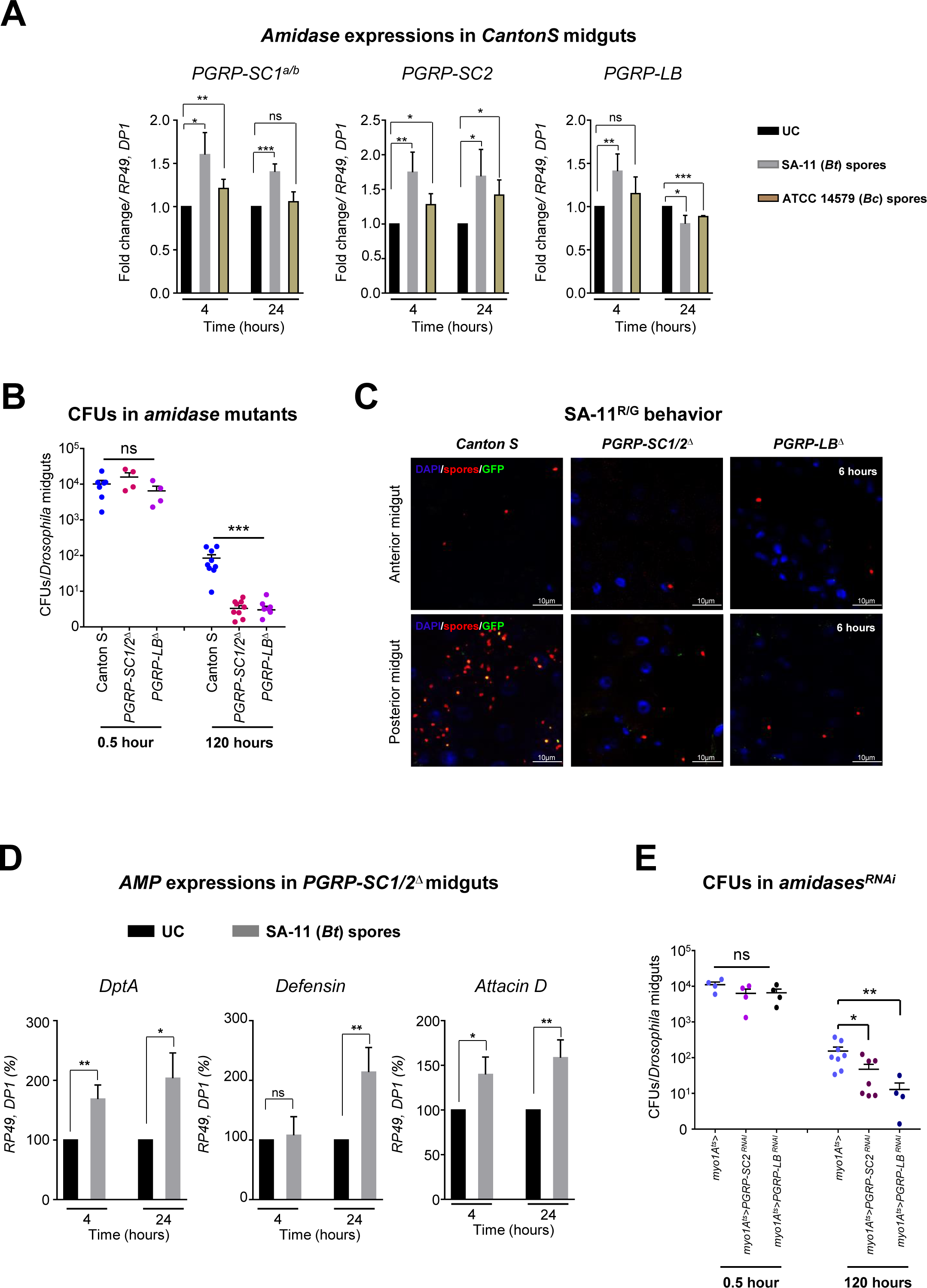
Amidases are involved in *Bt/Bc* persistence. **(A)** qRT-PCR analyses of *amidase* expressions in midguts upon SA-11 or *Bc* spore acute feeding. UC corresponds to flies fed with water. Results are shown as mean ± SEM of 10 flies from at least three independent experiments. **(B)** Bacterial load in midguts of *PGRP-SC1/2^Δ^* double mutant or *PGRP-LB^ΔE^* mutant 0.5 or 120 hours after SA-11 acute feeding with spores. **(C)** Representative confocal images showing SA-11^R/G^ spore germination in the anterior and posterior midgut of WT (Canton S), *PGRP-SC1/2^Δ^* double mutant or *PGRP-LB^ΔE^* mutant flies 6 hours after spore acute feeding. DAPI (blue) marks the nuclei. Spores are in red, vegetative cells in green. The yellow fluorescence corresponds to germinating spores (see Figure 2A). **(D)** qRT-PCR analyses of *AMP* expressions in midguts of *PGRP-SC1/2*^Δ^ mutants following acute feeding with SA-11 spores. UC corresponds to *PGRP-SC1/2*^Δ^ flies fed with water. mRNA levels in unchallenged *PGRP-SC1/2*^Δ^ flies were set to 100 and all other values were expressed as a percentage of this value. **(E)** SA-11 load in midguts of flies silenced for *PGRP-SC2* or *PGRP-LB* specifically in enterocytes (using the *myo1A^ts^* driver) 0.5 or 120 hours after acute feeding with spores. Student’s t-tests were used to analyze data in A and D. The Mann-Whitney test was used to analyze data in B and E. *p≤0.05, **p≤ 0.01, ***p≤0.001, ns = non-significant.

We therefore investigated whether the repression of *AMPs* expression upon spore ingestion was indeed dependent of amidases. As expected, expression of *AMPs* was not repressed; indeed, it was even induced by *Btk* SA-11 spore ingestion in a *PGRP-SC1/2^Δ^* or *PGRP-LB^Δ^* mutant background (Figure 4D compared with 3E). Because Amidases can be produced by the enterocytes or by the fat body (a systemic immune tissue) and can act at a distance from the site of production (Charroux et al., 2018), we specifically silenced *PGRP-SC2* or *PGRP-LB* in enterocytes and found a significant decrease in the load of *Btk* SA-11 in the midgut (Figure 4E). Overall, our data suggest that spores are not detected by the anterior midgut immune response (i.e., no production of ROS or AMPs), and reach the posterior regions where they germinate and can activate the Imd signaling target genes *PGRP-SC1*, *PGRP-SC2,* and *PGRP-LB*. In turn, Amidases promote a repression of basal expression of AMP-encoding genes. Consequently, downregulation of *AMP* expression favors *Bc/Btk* persistence in the posterior *Drosophila* midgut.

### The Imd pathway contributes to the spore intestinal persistence

Because the expression of *amidases* is under the control of the Imd pathway in the intestine, we first tested the involvement of the two Imd pathway receptors, PGRP-LC and −LE, in *Btk* SA-11 persistence. In flies homozygous viable for the loss-of-function mutant for either receptor, similar amounts of *Btk* SA-11 were ingested compared with WT flies after 30 minutes of spore feeding (Figure 5A). Nevertheless, 120 hours after feeding, only *PGRP-LE^112^* mutant flies showed a significant decrease in *Btk* SA-11 intestinal load (Figure 5A) similar to that observed for flies lacking the amidases (Figure 4B). These results show that PGRP-LE contributes to the sensing of germinated spores in the posterior midgut and confirm that PGRP-LE is the primary receptor in the posterior midgut that regulates amidase expression (Bosco-Drayon et al., 2012).

**Figure 5:**
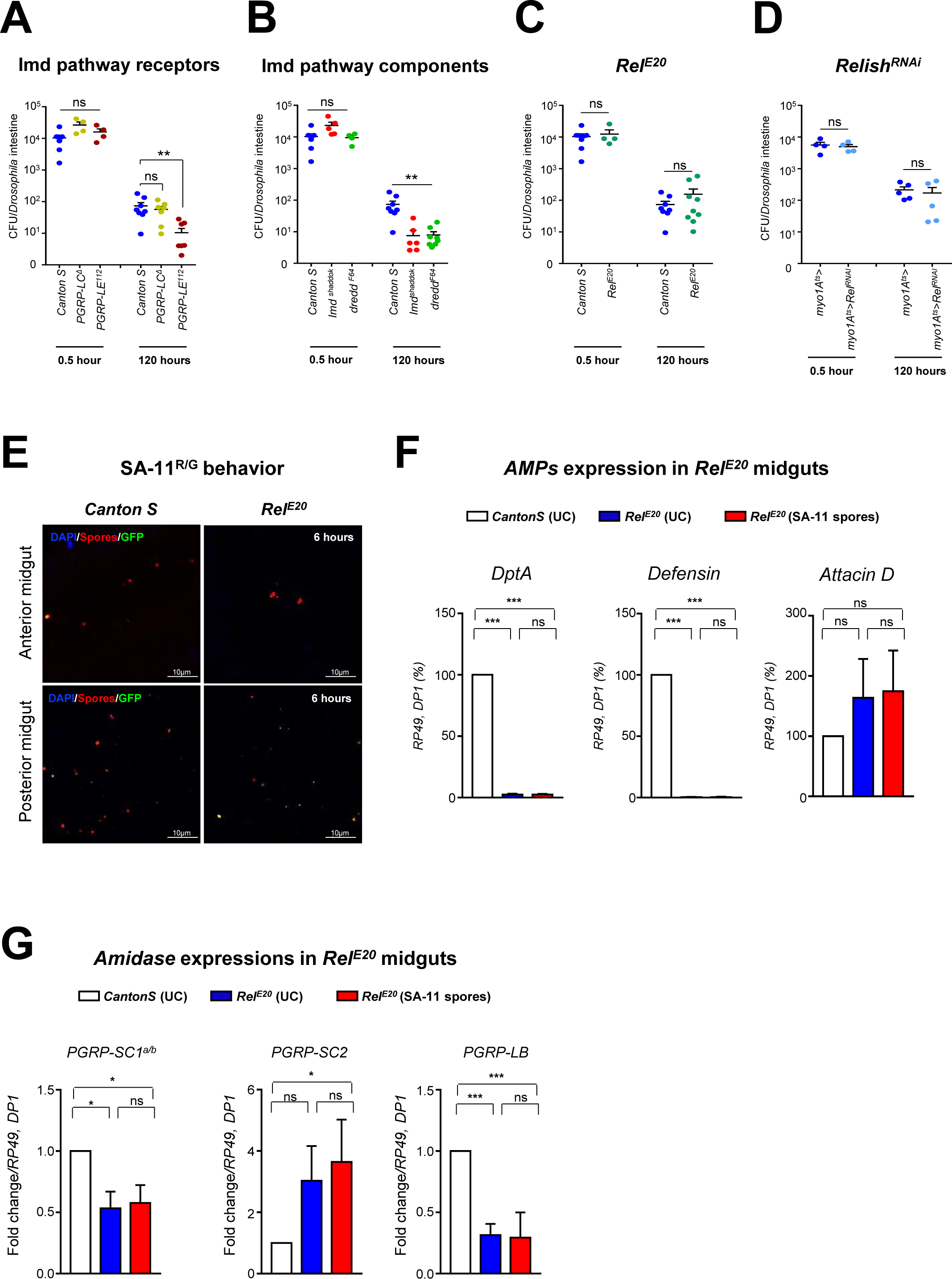
Imd pathway components but not the transcription factor (Relish) are involved in *Bacillus cereus* persistence. **(A-C)** SA-11 bacterial load in midguts of homozygous mutants for different components of the Imd pathway: *PGRP-LE^112^ and PGRP-LC^Δ^* (A), *Imd^shaddok^* and *Dredd^F64^* (B), *Rel^E20^* (C) 0.5 or 120 hours after acute feeding with spores. **(D)** SA-11 bacterial loads in midguts of flies silenced for *Relish* expression in the enterocytes (using the *myo1A^ts^-GAL4* driver) 0.5 or 120 hours after acute feeding with spores. **(E)** Representative confocal images showing SA-11^R/G^ spore germination in the anterior and posterior midgut of WT flies (Canton S) and *Rel^E20^* mutant flies 6 hours after acute feeding with spores. DAPI (blue) marks the nuclei. Spores are in red, vegetative cells in green. The yellow fluorescence corresponds to germinating spores (see Figure 2A). **(F and G)** RT-qPCR analyses of the expression of *AMPs* (F) and *amidases* (G) in *Rel^E20^* mutant flies 4 hours after acute feeding with SA-11 spores. UC corresponds to flies fed with water. Data represent mean ± SEM of at least four independent experiments. The Mann-Whitney test was used to analyze data in A-D. Student’s t-tests were used to analyze data in F and G. *p≤0.05, **p≤ 0.01, ***p≤0.001, ns = non-significant.

We further tested mutants for intracellular components of the Imd pathway. Loss-of-function mutants for the cytoplasmic components Imd (*imd^Shaddok^*) or Dredd (*Dredd^F64^*) also displayed a decrease in *Btk* SA-11 persistence 120 hours post-ingestion (Figure 5B). Unexpectedly, the bacterial load in flies homozygous mutant for the downstream Imd pathway NF-κB-like transcription factor Relish (*Rel^E20^*) (Hedengren et al., 1999) was similar to control flies (Figure 5C), although Rel has been found to be absolutely required in midgut epithelial cells to respond to enteropathogenic bacteria (Bosco-Drayon et al., 2012; Buchon et al., 2009b; Vodovar et al., 2005). To confirm the absence of *Rel* function in our model of spore infection, we inhibited *Relish* expression specifically in enterocytes. Silencing *Relish* in enterocytes did not change *Btk* bacterial abundance in the midgut (Figure 5D). In addition, we monitored the fate of the *Btk* SA-11*^R/G^* fluorescent strain in *Rel^E20^* mutants, six hours after acute ingestion. No obvious differences between *CantonS* and *Rel^E20^* were observed in the *Drosophila* midgut (Figure 5E), confirming that the transcription factor Relish had no significant role in the control of the germination and the persistence of *Btk* in the posterior midgut.

Given these observations, we wanted to test the involvement of Rel in controlling the remaining *AMP* expression in WT flies fed with spores (Figure 3E). We therefore performed qRT-PCR to measure the expression of the same *AMPs* in *Rel^E20^* mutant flies, either fed, or not, with spores. While *DptA* and *Defensin* were drastically downregulated in the absence of Rel at 4 hours (Figure 5F) and 24 hours (Figure S5A) in unchallenged flies or after spore ingestion, *AttD* expression was not affected in *Rel^E20^* flies, regardless of whether they were fed with spores (Figures 5F and S5A). In addition, the absence of Rel correlated with the downregulation of *PGRP-SC1a/b* and *PGRP-LB* expression (Figures 5G and S5B compared with 4A) but, unexpectedly, *PGRP-SC2* was strongly induced, even in the absence of spore ingestion (Figures 5G and S5B). The above data suggest that the expression of *PGRP-SC2* and, to a lesser extent, *AttD* in the posterior midgut are independent of the transcription factor Rel.

Taken together our data demonstrate that the Imd pathway can sense *Bc*/*Btk* geminating cells in the posterior midgut in a PGRP-LE-dependent manner in order to activate the expression of the amidases. In turn, amidases provoke a reduction of AMPs expression by tuning down the Imd pathway. Consequently, the decrease in AMP levels favor the local *Bc*/*Btk* persistence in the posterior midgut. Nevertheless, the most downstream component of the Imd pathway, Rel, appears not to be required.

### Dif cooperates with Relish to modulate the intestinal immune response to spore ingestion

These data prompted us to investigate the possible involvement of another NF-κB-related transcription factor in the *Drosophila* posterior midgut, the <u>D</u>orsal-related immunity factor (Dif), known to act downstream of the Toll pathway during the systemic immune response (Manfruelli et al., 1999; Meng et al., 1999; Petersen et al., 1995; Rosetto et al., 1995). To understand whether Dif could also be involved in *Bc/Btk* persistence, we took advantage of flies homozygous viable for the *Dif* loss-of-function allele, *Dif^1^*. While *Dif^1^* and WT flies ingested similar amounts of spores during the 30 minutes of feeding, 120 hours later, we observed a significant decrease of *Btk* SA-11 loads in *Dif^1^* mutant flies (Figure 6A). Confocal microscopy analysis confirmed the decrease of *Btk* SA-11*^R/G^* fluorescent cells, primarily in the posterior midguts of *Dif^1^* mutant flies, compared with WT (Figure 6B). To further analyze the role of Dif in the intestinal immune response, we monitored the expression of *AMPs* and *amidases* in *Dif^1^* mutant flies by qRT-PCR. In uninfected flies, the expression of *DptA* and *Defensin* was significantly lowered in the mutant compared with WT, while *AttD* was only slightly affected (Figures 6C and S6A). Feeding *Dif^1^* mutant flies with spores did not promote any induction or repression of *AMPs* (Figures 6C and S6A). Regarding the expression of *amidase*s, while they were all induced in a WT background upon spore ingestion (Figure 4A), in the absence of Dif, the expression of *PGRP-SC1* level was lower, while *PGRP-LB* remained inducible by the ingestion of spores (Figures 6D and S6B). Notably, contrary to what we observed in *Rel^E20^* mutants, *PGRP-SC2* was not upregulated in the absence of Dif (compare Figures 6D and S6B with Figures 5G and S5B).

**Figure 6.**
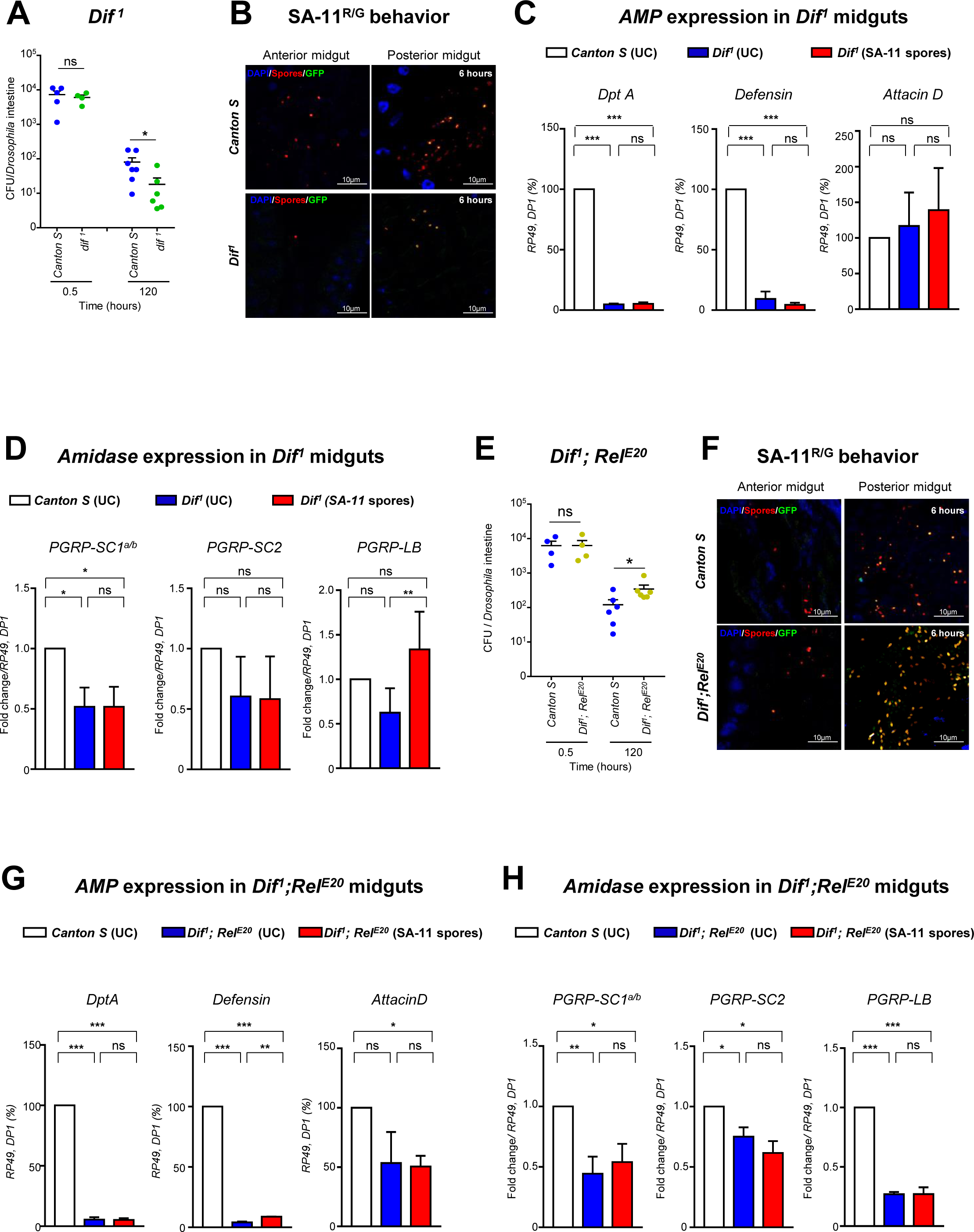
Dif and Relish are synergistically involved in *Bt* persistence. **(A and E)** SA-11 bacterial load in midguts of *Dif^1^* (A) or *Dif^1^;Rel^E20^* (E) homozygous mutants 0.5 or 120 hours after acute feeding with spores. **(B and F)** Representative confocal images showing SA-11^R/G^ spore germination in the anterior and posterior midgut of WT flies (Canton S) and *Dif^1^* (B) or *Dif^1^;Rel^E20^* (F) homozygous mutant flies 6 hours after acute feeding with spores. DAPI (blue) marks the nuclei. Spores are in red, vegetative cells in green. The yellow fluorescence corresponds to germinating spores (see Figure 2A). **(C, D, G and H)** RT-qPCR analyses of the expression of *AMPs* (C and G) and *amidases* (D and H) in *Dif^1^* (C and D) or *Dif^1^;Rel^E20^* (G and H) homozygous mutant flies 4 hours after acute feeding with SA-11 spores. UC corresponds to flies fed with water. Data represent mean ± SEM of at least four independent experiments. The Mann-Whitney test was used to analyze data in A and E. Student’s t-tests were used to analyze data in C, D, G and H. *p≤0.05, **p≤ 0.01, ***p≤0.001, ns = non-significant.

To confirm the cooperation of both NF-κB factors in regulating the expression of *amidases* and *AMPs* in the posterior midgut, we generated a *Dif^1^; Rel^E20^* double mutant flies. We first monitored *Btk* persistence upon spore ingestion in this genetic background. *Dif^1^; Rel^E20^* double mutant and WT flies ingested a similar amount of spores during the 30 minutes of feeding (Figure 6E). However, 120 hours later there was a significantly higher *Btk* SA-11 load in the double mutant flies compared with WT (Figure 6E). Consistently, 6 hours after ingestion of *Btk* SA-11*^R/G^* spores, more fluorescent bacteria were present in the posterior midguts of *Dif^1^; Rel^E20^* double mutant flies compared with WT (Figure 6F). In this double mutant background, the expressions of *AMPs* and *amidases* were lowered (Figures 6G, 6H, S6C and S6D).

Together our data suggest that Dif and Relish cooperate to regulate the expression of *DptA*, *Defensin* and *PGRP-SC1*, while Dif and Rel likely act redundantly to regulate the expression of *AttD*. Moreover, Relish and Dif have opposite effects on *PGRPSC-2* expression: Relish represses *PGRPSC-2* while Dif activates it. Finally, *PGRP-LB* expression appears to be regulated only by Relish. Overall, these results demonstrate that Dif and Relish cooperate to tightly balance the expression of AMPs and amidases in the posterior midgut of unchallenged as well spore-fed flies.

### *Bacillus* spore persistence leads to *amidase* expressions in the mouse gut

Because the *Bc* group is involved in human digestive illnesses, we wondered whether their spores could also persist in the intestine of mammals. We, therefore, investigated the fate and behavior of spores in the mice. In *Drosophila* midguts, all the spores we have studied behave similarly (Figures 1, S1, 2 and S2), therefore, to ethical considerations in mouse experiments, we restricted our experiments to the *Btk* SA-11 strain. Mice were fed with vegetative cells or spores using oral gavage (Figure 7A). In mammals, the small intestine is divided into three functional sections: the duodenum, the jejunum, and the ileum. Despite the duodenum being smaller than the other two more posterior regions of the small intestine, we decided to split the small intestine into three equivalent domains since there is no obvious anatomical distinction (Figure S7A).

**Figure 7.**
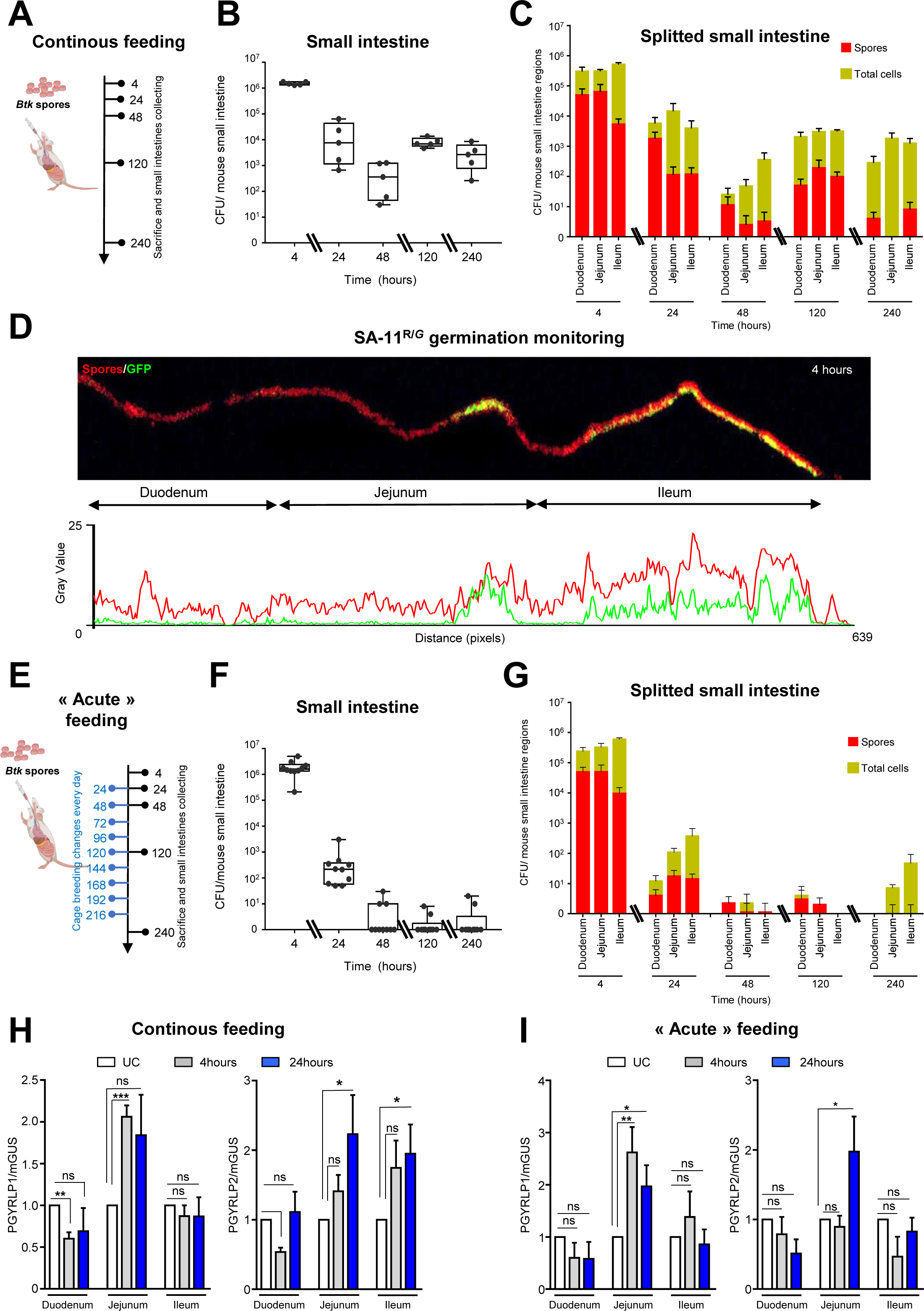
*Bacillus* spore persistence leads to *amidase* expressions in the mouse gut. **(A)** Schematic illustration of the procedure for oral gavage using SA-11 spores. Mice were kept in the same cage throughout the experiment. Small intestine samples were collected at tC=C4, 24, 48, 120 and 240 hours after the gavage. **(B and F)** SA-11 bacterial loads in split mouse small intestines after gavage using the mode of breeding shown in (A) for (B) and in (E) for (F). The vertical axis indicates the median number of CFUs per mouse. **(C and G)** SA-11 bacterial load in different parts of the small intestines using the mode of breeding shown in (A) for (C) and in (E) for (G). Green bars (non-heated samples) represent the whole SA-11 bacterial load (spores and vegetative cells). Red bars (heated samples) represent the proportion of non-germinated spore loads. Data represent the mean ± SEM of at least five independent experiments. **(D)** Monitoring of SA-11^R/G^ spore germination in the small intestine 4h post-gavage. The lower panel represents plots of the average fluorescence intensity (the mean gray value) of SA-11^R/G^ measured along the small intestine in the upper panel. Red labeling corresponds to spores and green (GFP) corresponds to vegetative cells (GFP). **(E)** Schematic illustration of the procedure for acute oral gavage using SA-11 *Bt* spore. Small intestines samples were collected at tC=C4, 24, 48, 120 and 240 hours after the oral gavage. Breeding cage was changed initially 4 hours post-oral-gavage and then every day. **(H and I)** RT-qPCR analyses of the expression of mouse amidases (*Pgylrp1* and *Pgyrlp2*) in different parts of the small intestine after spore ingestion using the mode of breeding shown in (A) for (H) and in (E) for (I). Error bars represent SEM (n=5). The student’s t test was used to analyze statistical significance. *p≤0.05, **p≤ 0.01, ***p≤0.001, ns = non-significant.

As expected, we first found that upon gavage with vegetative cells, *Btk* SA-11 were rapidly cleared from the small intestines of mice (Figures S7B and S7C) within a time frame similar to that of *Drosophila*. By contrast, gavage with spores led to the detection of *Btk* SA-11 in the small intestine (equivalent to the *Drosophila* midgut) 240 hours post-oral-gavage (Figure 7B) as in *Drosophila* midguts. To monitor the spore germination, we performed CFU measurements in heat-treated *vs* non-heat-treated intestinal samples. We found that 4 hours post-gavage, *Btk* SA-11 spores were evenly distributed all along the small intestine and that vegetative cells were more abundant in the ileum (Figure 7C). The few vegetative cells we detected in the duodenum could be explained by the 1/3 splitting method we used that encompassed the whole duodenum and likely part of the anterior jejunum (Figure S7A). We also observed an obvious *Btk* SA-11 germination localized in the jejunum and ileum, 24 hours onwards post-gavage (Figure 7C). Next, to confirm the timing and locations of spore germination in the mouse intestine, we forced-fed our mice with the *Btk* SA-11*^R/G^* strain. Two hours post-gavage, *Btk* SA-11*^R/G^* spores were localized throughout the small intestine and few vegetative cells were detectable in the posterior ileum (Figure S7D). Four hours post-gavage, *Btk* SA-11*^R/G^* vegetative cells occupied the posterior part of the jejunum and the ileum (Figure 7D). Twenty-four hours post-gavage, we detected mostly vegetative cells throughout the jejunum and the anterior part of the ileum (Figure S7E). Notably, spores were evenly distributed along the small intestine at the different time points analyzed (Figures 7D and S7D and S7E). Together, these results suggested that spores persist and germinate in the mammalian posterior small intestine regions, similar to our observations in the *Drosophila* midgut.

However, as rodents are known for coprophagy, the data obtained above might be due to continuous ingestion of spores beyond initial gavage (Kenagy and Hoyt, 1979). Consistent with this, we detected *Btk* SA-11 spores in fecal samples up to ten days post-gavage (Figure S7F). To prevent this, we frequently changed breeding cages (Figure 7E). As shown in (Figure 7F), *Btk* SA-11 bacteria were detectable 24 hours post-gavage in the small intestine of all mice (n=10). Interestingly, two mice still harbored *Btk* SA-11 bacteria 240 hours post-oral-gavage by spores. We then monitored spore germination using CFU measurements in heat-treated *vs* non-heat-treated intestinal samples. Four hours post-gavage, *Btk* SA-11 spores were evenly distributed all along the small intestine, and vegetative cells were more abundant in the ileum (Figure 7G). Twenty-four hours post gavage, we also observed *Btk* SA-11 germination preferentially localized in the jejunum and ileum (Figure 7G). Later, *Btk* SA-11 spores were still detectable in the duodenum, while vegetative cells were barely detectable. Ten days after gavage, only vegetative cells were detected in the jejunum and ileum (Figure 7G). Collectively, our results demonstrate that spores can persist for several days in the mouse small intestine and germinate preferentially in the posterior part of the small intestine, the jejunum and the ileum, similar to our observations in *Drosophila* midguts.

Finally, we explored the possibility that the two mouse orthologs of the *Drosophila* amidases Pglyrp-1 and Pglyrp-2 (Lee et al., 2012; Osanai et al., 2011), might be upregulated in response to spore ingestion. The expression of *Pglyrp-1* and *Pglyrp-2* genes was monitored in the murine small intestine by quantitative RT-qPCR upon gavage with *Btk* SA-11 spores, with or without changing the breeding cages (Figures 7A and 7E). We observed, 4-and 24-hours post-gavage, an upregulation of *Pglyrp-1* only in the jejunum (Figure 7H and 7I), whereas *Pglyrp-2* was upregulated both in the jejunum and ileum without changing the breeding cage (Figure 7H) and only in the jejunum upon frequent breeding cage change e to avoid continuous consumption (Figure 7I). These results demonstrate that spores can persist in the small intestine of mice, germinate in the posterior parts, and trigger the expression of amidases. Altogether our data suggest an evolutionarily conserved role for amidases in the innate immune response and in tolerance to bacterial spores.

## DISCUSSION

The majority of *Bc*-dependent FBOs, is due to the ingestion of *Bc* bacteria, which must grow in the gut and subsequently produce pore-forming enterotoxins responsible for the onset of diarrhea symptoms (Jovanovic et al., 2021). However, the mechanisms by which *Bc* bacteria colonize the gut and produce toxins remain poorly understood, and several questions unanswered. Is the disease due to the ingestion of vegetative bacteria or spores? What is the distribution and the fate of ingested spores along the gastrointestinal tract? How does the intestinal innate immune system detect and fight the infection? Here, we deciphered the behavior and fate of *Bc* cells in the intestine of both *Drosophila melanogaster* and mouse, and demonstrated that spores escape the innate immune system to reach the posterior part of the midgut/small intestine, where they can germinate and persist for days within the animal.

First of all, our findings confirm *in vivo* that after ingestion of vegetative cells, the *Bc* load in the *Drosophila* and mouse intestine remains low and that the bacteria are cleared in less than 24 hours (Benguettat et al., 2018; Loudhaief et al., 2017; Rolny et al., 2014). In *Drosophila*, it has also been shown that the presence of vegetative cells in the anterior midgut is rapidly detected, triggering the production of ROS and visceral spasms, both cooperating to quickly evict the undesired bacteria (Benguettat et al., 2018). Therefore, the minimum infectious dose required to cause intestinal disorders (at least 10^5^ CFU, (Berthold-Pluta et al., 2015)) is likely difficult to reach upon ingestion of vegetative cells. On the contrary, it has been suggested that the capacity of spores to withstand extreme conditions would allow them to overcome stomach acidity and bile salt attacks in the duodenum, favoring germination in the posterior small intestine. Consequently, the infectious dose could be more readily achievable, as illustrated by the 10^3^ spores/g of food that can be associated with FBOs (Bonis et al., 2021; EFSA BIOHAZ, 2016). Importantly, our *in vivo* data reveal the conserved fate and behavior of spores in the intestine of *Drosophila* and mice. The spores of all *Bc* group strains we tested persisted up to 10 days post-ingestion. Consistent with our observations, studies have shown that *Bc* can persist at least 18 days in the intestine of rats transplanted with human-flora (Wilcks et al., 2008) and 30 days in mouse intestine (Oliveira-Filho et al., 2009). Moreover *Bt* could be detected in fecal samples in greenhouse workers five days after cessation of bioinsecticide use (Jensen et al., 2002). Interestingly, we also showed *in vivo* that spores accumulate and germinate in the posterior part of the *Drosophila* midgut and mouse small intestine. Similarly, data suggest that spores derived from the probiotic *B. subtilis* germinate in the jejunum and eventually in the ileum of mice (Casula and Cutting, 2002; Tam et al., 2006). Our data also highlight the very rapid germination of spores in the posterior parts of the intestine (in less than 2 hours). Indeed, while the proximal regions of the *Drosophila* midgut and mouse small intestine are quite acidic and produce digestive enzymes to break down food, the more distal parts of the intestine harbor a more basic and anaerobic environment with nutrient availability (Marianes and Spradling, 2013; Osman et al., 2012; Shanbhag and Tripathi, 2009). Interestingly, it has been shown, *in vitro*, that anaerobic conditions slow down the growth rate of *Bc* but favor the production of CytK, Nhe and Hbl enterotoxins (Berthold-Pluta et al., 2015; Duport et al., 2004; Duport et al., 2006; Jovanovic et al., 2021; Miller et al., 2021; van der Voort and Abee, 2009; Zigha et al., 2006). Hence, all the conditions for spore germination and enterotoxin production are encountered in the posterior *Drosophila* midgut and mouse small intestine, which accounts for the occurrence of diarrheic symptoms when a critical bacterial load is reached.

Importantly, our data also show that the local innate immune response is ineffective in eliminating vegetative cells in the posterior regions of the *Drosophila* midgut or mouse small intestine, which enables *Bc/Bt* persistence. Using *Drosophila* genetic tools, we first show that spores are not detected in the anterior midgut, unlike vegetative cells, which rapidly trigger immune ROS production (Benguettat et al., 2018; Lee et al., 2013). Strikingly, although spores germinate in the posterior midgut, there is no release of immune ROS. Immune ROS are normally produced in a DUOX-dependent manner in response to uracil secretion by allochthonous bacteria (Lee et al., 2013). Uracil is thought to serve as bacterial growth factor, promoting proliferation (Du et al., 2016). Hence, we can assume that either *Bc/Bt* vegetative scells in the posterior midgut do not produce uracil or the host receptor for uracil (Lee et al., 2015) is absent from the posterior midgut. Moreover, the germination of spores in the posterior midgut, through the activation of the negative regulators, amidases, dampens the production of AMPs. Consequently, the combination of the absence of ROS and the reduced levels of AMPs favor *Bc/Bt* persistence.

Why does spore germination induce only the genes encoding amidases and not those encoding AMPs in the posterior midgut? It has been shown that the Imd pathway cytosolic receptor PGRP-LE is required all along the fly midgut to activate *AMP* genes in the anterior in response to pathogenic bacteria and to upregulate the amidases PGRP-SC1 and −LB in the posterior midgut in response to commensal bacteria. The transmembrane PGRP-LC receptor is also required in cooperation with PGRP-LE in the anterior midgut to activate the expression of *AMPs* in response to pathogenic bacteria, however, PGRP-LC is dispensable in the posterior midgut (Bosco-Drayon et al., 2012; Buchon et al., 2013; Costechareyre et al., 2016; Neyen et al., 2012; Zhai et al., 2018a). Consistent with this observation, we found that only PGRP-LE is involved in response to spore ingestion. Therefore, in the posterior midgut, the germination of spores allows the activation of the Imd pathway in a PGRP-LE dependent manner, leading to the induction of *amidases* but not of *AMPs*. Hence the germination of spores of *Bc/Bt* in the posterior compartment are perceived as if they were commensal bacteria, inducing a tolerance response (Bonnay et al., 2013; Bosco-Drayon et al., 2012; Morris et al., 2016) through the induction of Amidases that in turn dampen *AMPs* expression. Our spore ingestion paradigm also reveals that when the Imd pathway is only mobilized in the posterior midgut (spores are not detected in the anterior compartment) in a PGRP-LE-dependent manner, the only response elicited is the induction of amidases, even if the ingested bacteria are non-commensal ones. Consistently, the transcription repressor Caudal has been shown to be involved in the repression of *AMP* expression specifically in the posterior midgut (Ryu et al., 2008). Hence the midgut could be separated into two distinct immune domains: the anterior midgut is competent to fight pathogenic bacteria ingested along with the food, and the posterior midgut is immune-tolerant to sustain commensal flora. *Bc/Bt* spores have developed a strategy to hijack this physiological state for their own benefit, allowing them to escape the strong anterior immune response that would otherwise kill the germinated cells. Consistent with this model, it has been well demonstrated that the *Drosophila* posterior midgut is capable of increased cell turnover, when compared to the anterior midgut, in order to overcome the damages caused by pathogens (Apidianakis et al., 2009; Jiang et al., 2009; Marianes and Spradling, 2013; Tamamouna et al., 2020; Zhou et al., 2013), likely to compensate for a weaker innate immune response. Interestingly, our data show that spores behave similarly in the mouse small intestine, germinating preferentially in the jejunum and ileum. Moreover, the two mouse orthologs of the *Drosophila* amidases, *Pglyrp1* and *Pglyrp2* (Lee et al., 2012; Osanai et al., 2011) are also induced in the jejunum and ileum, suggesting a conserved intestinal physiological response to spore ingestion.

Unexpectedly, our work in *Drosophila* also revealed that Relish is not the sole NF-κB factor driving the innate immune response in the posterior midgut. Indeed it has been well demonstrated that Relish is absolutely required in the anterior midgut downstream of the Imd pathway to mount an efficient immune response against pathogens (Bosco-Drayon et al., 2012; Buchon et al., 2009a; Buchon et al., 2009b; Cronin et al., 2009). However, we show that in the posterior midgut, Dif intervenes to control *DptA* and *Defensin* activation, probably in cooperation with Relish, since the absence of one of the NF-κB factors is sufficient to shut down their expression. In agreement, it has been shown that during the systemic immune response, both NF-κB factors were able to form hetero-and homodimers to differentially control *AMP* genes (Han and Ip, 1999; Morris et al., 2016; Tanji et al., 2010). Similarly, Relish/Dif heterodimers likely regulate *PGRP-SC1*, since its expression is lost in either *Dif* or *Rel* loss-of-function mutants. However, induction of *PGRP-LB* appears to be only under the control of Relish. Finally, Dif and Relish exert opposite effects on *PGRP-SC2* expression. While Rel represses its expression, Dif is required for its induction. Interestingly, it has been shown that the IκB factor Pickle can bind to Relish homodimers, converting them into transcriptional repressors of *AttD* expression (Morris et al., 2016). Therefore, a combination of NF-κB homo-and hetero-dimers, plus the presence of specific negative regulators, fine-tune the posterior immune response, limiting the level of expression of *AMPs* to enable commensal flora to become established, but also unfortunately allowing some opportunistic bacteria to persist. Interestingly, a role for Dif in shaping the intestinal commensal flora, downstream of the Toll pathway, has recently been uncovered (Bahuguna et al., 2022). Along with our results, this suggests that the Toll pathway could also be active in the posterior midgut and may contribute to the immune response against pathogens.

Together, our data shed light on the conserved behavior and strategy of *Bc/Bt* spores to escape the innate immune response in the proximal part of the small intestine, allowing them to reach and germinate in the distal region. Our work also provides useful tools for further investigation to understand when and how enterotoxins are produced and trigger diarrheic symptoms. Our work also highlights that the persistence and load of *Bc/Bt* can be enhanced and could potentially lead to more severe symptoms in immunocompromised individuals.

## AKNOWLEDGEMENT

We would like to thank Bernard Charroux, François Leulier, Julien Royet, Heinrich Jasper, Leopold Kurz, Mark Hanson and Bruno Lemaitre for kindly providing fly stocks. We also thank Didier Lereclus for providing the pHT315 plasmid. We are grateful to Leanne Jones, Raphaël Rousset, Carmelo Luci, Bernard Charroux and Bruno Lemaitre for their advice on the project and the manuscript. We also thank Olivier Pierre and Anne Doye for their help for the microscopy, Dorota Czerucka for mouse intestine dissection, and Olivia Benguettat for her technical help. We greatly acknowledge the C3M Animal core facility. Our thanks to the Université Côte d’Azur Office of International Scientific Visibility for English language editing of the manuscript. We also thank the Space, Environment, Risk and Resilience Academy of the Université Côte d’Azur for their financial support.

## AUTHOR CONTRIBUTIONS

Conceptualization: S.H., L.B. and A.G.

Methodology: S.H., A.B.-B., P.M., M.-P.N.-E., G.M. and M.B.

Validation: S.H., A.B.-B., P.M., A.F., M.B., L.B. and A.G.

Formal Analysis: S.H., A.B.-B., P.M., A.F., L.B. and A.G.

Investigation: S.H., A.B.-B., P.M., M.-P.N.-E. and M.B.

Data Curation: S.H., A.B.-B., P.M., A.F., M.B. and A.G.

Writing – Original Draft: S.H. and A.G.

Writing – Review & Editing: S.H., A.B.-B., M.B., L.B. and A.G. Visualization: S.H., L.B and A.G.

Supervision: L.B and A.G.

Project Administration: L.B. and A.G.

Funding Acquisition: M.B., R.R., L.B. and A.G.

## EXPERIMENTAL PROCEDURES

### Bacterial strains

The two bioinsecticide strains (SA-11 and ABTS-351) were used as commercial formulations. In parallel, the strain ABTS-351 was also used after bacterial isolation and “home-made” spore production as described below. The *Btk^ΔCry^* strain (#4D22) was collected from the Bacillus Genetics Stock Center (www.bgsc.org) (González et al. 1982). The *Bc* (#ATCC 14579) was provided by ANSES Maisons-Alfort. *B. toyonensis* strain were selected in this work. *Erwinia carotovora carotovora 15* (*Ecc 15)* was kindly provided by Bruno Lemaitre (Basset et al., 2000). Bacterial strains were grown in LB medium at 30°C for 16 h.

### Construction of SA-11**^Δ^***^Cry^*

The mutant SA-11*^ΔCry^* was obtained from the WT strain SA-11, by a procedure of plasmid curing, as follows. After isolation on TSA-YE agar (Biomérieux, 18h culture at 30°C), the strain SA-11 was sub-cultured successively 3 times in 10ml of brain-heart Infusion (BHI, Oxoid) broth at 42°C with agitation, for 64, 48 and 36h respectively. The first BHI culture was inoculated from an isolated colony, and the subsequent cultures were inoculated with 100µl of the previous ones. Clones from the last culture were isolated on TSA-YE agar after serial dilution, then subcultured on the sporulating medium hydrolysate of casein tryptone (HCT) + 0.3% Glc, to select clones unable to produce crystals visible by phase contrast microscopy. The absence of plasmids carrying the *cry* genes was checked by sequencing. Briefly, the genomic DNA of SA-11*^ΔCry^* and SA-11 WT were extracted using the KingFisher cell and Tissue DNA kit (ThermoFisher) and sequenced with Illumina technology at the Institut du Cerveau et de la Moelle Epinière (ICM) platform, as previously described (Bonis et al., 2021), (SAMN23436137 and SAMN23455549, respectively). The absence of cry genes in SA-11*^ΔCry^* has been confirmed from raw reads, using KMA (Clausen et al., 2018). Consistently, a plasmid reconstruction made with Mob-Suite (Robertson and Nash, 2018) suggested the loss of 4 plasmids in SA-11*^ΔCry^* compared with SA-11 WT.

### Btk^ΔCry-GFP^, SA-11^GFP^, SA-11^ΔCryGFP^, Bc^GFP^ and B. toyonensis^GFP^ strains

The GFP coding sequence was inserted into the *pHT315* plasmid (bearing the erythromycin-resistant gene) (Theoduloz et al., 2003) (gift from Didier Lereclus). The *pHT315-GFP* recombinant plasmid was transfected and amplified into competent *dam^-^/dcm^-^ E. Coli* (NEB#C2529H) which allowed it to be demethylated. *pHT315-GFP* was then extracted and purified using either the Pureyield plasmid miniprep kit (Promega #A1223) or the QIAGEN® Plasmid Mini Kit (QIAGEN). For the extraction using the QIAGEN® Plasmid mini Kit, the DNA solution was concentrated by isopropanol precipitation following the manufacturer’s recommendations and resuspended in PCR-grade water. The DNA concentrations were measured using the NanoDrop1000 spectrophotometer (Thermo Fisher Scientific).

The different strains from the *Bc* group were transfected with the *pHT315-GFP* plasmid as follows. Strains were plated on TSA-YE agar at room temperature for 48h, then subcultured in 10 ml of BHI for 18h at 30°C, after inoculation from isolated colonies. The cultures were diluted 1/100 in 100 ml BHI and incubated at 37°C under agitation until an OD600_nm_ of about 0.3 was reached. Bacteria were washed in 10 ml of cold electroporation buffer (400 mM sucrose, 1mM MgCl2, phosphate-buffered saline 1X, pH 6.8) and then resuspended in 850µl of cold electroporation buffer. A hundred µl of each suspension was incubated with 250 ng of plasmid DNA in ice for 5 min, then submitted to electroporation using the MicroPulser Electroporator (Biorad, program Sta), and 2 mm electroporation cuvettes. After the addition of 0.9 ml of BHI, bacteria were incubated for 2h at 37°C and isolated on TSA-YE agar supplemented with 10 µg/ml of erythromycin. The selected clones were checked for the presence of the *GFP* gene after sequencing (SAMN23436138 and SAMN23436139, respectively), and for the expression of GFP using fluorescence microscopy.

### Spore production

Strains were plated on LB-agar plates and grown overnight at 30◦C. Bacteria were grown at 30°C in HCT-agar medium (pH 7.2) containing per 1L: 5g tryptone (Oxoid), 2g casein hydrolysate (Oxoid), 15g agar, 3g glucose, and salts as previously reported in a sporulation-specific medium (Lecadet et al., 1980). After 10 days of incubation, spores were washed with 0.15% NaCl and heat-treated for 20 min at 70°C. Then cells were centrifuged at 10000g, 8°C for 20min. Spores were washed with sterile deionized water and centrifuged at 10000g, 8°C for 20min. The supernatant was discarded, and the washing was repeated once. The last pellets were taken up in 10ml and lyophilized (freeze-drying equipment model: RP2V). The numbers of spores produced were determined by estimating the CFUs on LB plates after serial dilution of lyophilized spores.

### Time lapse fluorescence of **SA-11***^R/G^* spore germination

SA-11*^R/G^* spores were placed on 1.5% agarose pads on a microscopy slide and covered with a cover glass. The use of agarose pad allowed for stabilizing spores to be achieved in the microscopy samples. The agarose pads were incubated at 37°C for 60min to accelerate spore germination process. The time lapse images were taken once every 5 minutes for 90 minutes to avoid bleaching. Images were acquired using the Zeiss LSM 880 microscope equipped with the AiryScan detector.

### *Drosophila* rearing and stocks

Flies were reared on our standard laboratory medium (Nawrot-Esposito et al., 2020) in 12h/12h light/dark cycle-controlled incubators. We used the following stocks: WT *canton S* (Bloomington #64349); *W^1118^* (Bloomington #3605); *Rel^E20^* (Bloomington #55714); *w; PGRP-LC*^Δ*E*^ (Bloomington #55713); *yw; PGRP-LE^112^* (Bloomington #33055); *w; PGRP-SC*^Δ^ (Bloomington #55724); w; *PGRP-LB*^Δ^ (Bloomington #55715; gift from B. Charroux); *Dredd^F64^* (Leulier et al., 2000) (gift from B. Charroux); *imd^Shaddok^* (gift from B. Charroux *); w; Dif^]^* (Bloomington #36559); Δ*AMP14* (gift from B. Lemaitre) (Carboni et al., 2022); *w; AttD-Gal4 UAS-cherry* (gift from Leopold Kurz) (Tavignot et al., 2017)*; w; DptA-cherry* (gift from Leopold Kurz, Bloomington #55706); *UAS-DUOX^RNAI^* (Bloomington #38907); *UAS-REL^RNAI^* (Bloomington #33661); *UAS-PGRP-LB^RNAI^* (Bloomington #67236); *UAS-PGRP-SC2^RNAI^* (Bloomington #56915); *w; myo1A-Gal4; tubGal80ts UAS-GFP/TM6b* (gift from Nicolas Tapon) (Shaw et al., 2010). The *w; Rel^E20^, Dif^1^* homozygous viable double mutant was obtained using classic mendelian genetic crosses.

### *Drosophila* oral intoxication

Five-six-days old virgin females *Drosophila* were reared at 25°C. For conditional expression of UAS-GAL80^ts^linked transgenes, flies were developed at room temperature, then shifted to 29°C for 7 days to induce transgene expression. Before intoxication, *Drosophila* females were put into a new vial without medium for starvation for 2 hours at 25°C or at 29°C for UAS-GAL80^ts^ flies. This allows the synchronization of food intake once in contact with the contaminated medium. Ten females were transferred into a *Drosophila* narrow vial containing fly medium covered with a filter disk where the spore solution was deposited. The inoculum used for continuous and acute intoxication were respectively 10^6^ CFU/5 cm^2^/fly and 10^8^ CFU/5 cm^2^/fly respectively. For the acute intoxication, *Drosophila* were fed for 30 minutes with the spore inoculum, then transferred to a new sterile vial until dissection. For the continuous intoxication, *Drosophila* were let in contact with the spore inoculum until the dissection time. For control conditions, *Drosophila* females were fed with sterile deionized water in the same conditions.

### Bacterial load quantification (CFU) in *Drosophila* midgut

Flies were washed first in ethanol 70% and then in PBS1X before guts dissection in PBS1X. Whole midguts or split parts (anterior and posterior regions) were crushed in 200µL of LB at various times post-ingestion using a micro pestle and the homogenate was serially diluted in LB and incubated overnight at 30°C on LB agar plates. Colony counting was performed the day after.

### Bacterial load quantification (CFU) on the filter disk

The filter disk was washed and vortexed in 1ml of sterile water. The homogenate was serially diluted in sterile water and incubated overnight at 30°C on LB agar plates. Colony counting was performed the day after.

### Heat-treatment

The intestinal samples or the filter disk samples were heated at 75°C for 25 min to kill the germinating spores and the vegetative cells. Afterward, the spores were enumerated as described above.

### *In vivo* monitoring of spore germination

Flies were fed with *Btk* SA-11^R/G^. Guts were dissected and fixed with 4% formaldehyde in PBS1X for 20min and immediately mounted in Fluoroshield-DAPI medium. Images acquisition was performed at the microscopy platform of the Institut Sophia Agrobiotech (INRAE 1355-UCA-CNRS 7254-Sophia Antipolis) with the microscope Zeiss AxioZoom V16 with an Apotome 2 and a Zeiss LSM 880 microscope equipped with the AiryScan detector. Images were analyzed using ZEN and Photoshop software and ImageJ.

### RNA extraction and Real-time qPCR for *Drosophila* guts

Four biological replicates were generated for each condition. Total RNA was extracted from 10 *Drosophila* midguts using Microelute Total RNA kit (Omega Bio Tek) and dissolved in 20µl of RNase-free water. The quantity and quality of RNA were assessed using a Thermo Scientific™ NanoDrop 2000. 550ng of extracted RNA was reverse transcribed to cDNA using Qscript™. Real-time PCR was performed on AriaMX Real-Time (Agilent) in a final volume of 20µl, using the EvaGreen kit. Each experiment was independently repeated three times. Relative expression data were normalized to *RP49* and *Dp1* genes

### HOCl staining with R19S

The protocol is described in (Hachfi et al., 2019).

### The ethics statement for mouse model

This study was carried out in strict accordance with the guidelines of the Council of the European Union (Directive 86/609/EEC) regarding the protection of animals used for experimental and other scientific purposes. The protocol was approved by the Institutional Animal Care and Use Committee on the Ethics of Animal Experiments of Nice, France (APAFIS#18923-2019012512125055 v3).

### Mice

Female Balb/c mice (7 weeks of age) were used. The mice were matched by age with a control group. The mice were kept in animal house facilities with 12:12-h light/dark cycles in standard animal cages and fed with a standard pellet diet, as well as a plastic bottle. Mice were acclimatized to these conditions for 1 week before entering the study.

### Mice oral administration

Balb/c mice (7 weeks, female) were divided into two groups (n=5-10). There was no significant difference in body weight between the groups. Mice received 200µl of bacterial suspension or sterile PBS1X. The bacterial suspension was prepared by dissolving spores/bacterial at a concentration of 10^8^ CFU/mouse. All mice were transferred to freshly sterilized cages every day.

### Bacterial load quantification (CFU) in mice small intestines

At the time of dissection, Mice received freshly prepared Ketamine-Xylazine (10mg/ml)- 10mg/ml) mixture doses. The anesthetic mixture was injected intraperitoneal IP. Blood was collected from a tail vein sampling. Then, mice were euthanized. Duodenum, jejunum, ileum were sampled, weighed, and crushed in 1ml of sterile PBS1X using the precellys lysing kits CK14 (Bertin Technologies) The homogenate was serially diluted in PBS1Xx and incubated overnight at 30°C on LB agar plates. Colony counting was performed the day after.

### *In-vivo* monitoring of spore germination in mice small intestines

Mice fasted for 48h SA-11*^GFP^*spores suspension per gavage at a concentration of 10^8^ CFU/mouse. Guts were dissected and immediately visualized with PhotonIMAGER Optima (Biospace Lab, Nesles-la-VallCe, France). The data was analysed using M3 Vision software (Biospace Lab, Nesles-la-VallCe, France) version 1.1.2.26170 and Fuji software.

### RNA extraction and RT-qPCR for mice guts

One centimeter of intestinal tissue (duodenum, jejunum, or ileum) was homogenized in 2mL tubes containing 300µl Trizol using the Precellys® system. The duodenum part was taken directly after the stomach, the jejunum part was taken in the middle, and the ileum was the distal part near the cecum. After extraction with chloroform, precipitation with isopropanol and washings with 70% ethanol, and extracted RNA was resuspended in 50μL of sterile distilled water. cDNA synthesis was performed with Applied Biosystems™ Kit (Thermo Fisher Scientific). Gene expression was analyzed by quantitative reverse transcription PCR (qRT-PCR) analysis on a QuantStudio™ 5 Real-Time PCR System (Thermo Fisher Scientific) and normalized to endogenous control gene mGUS. Gene expression is depicted as relative mRNA amounts (relative quantities (RQ) after normalization to the expression of endogenous control gene mGUS calculated using delta/delta Ct method with the software provided by QuantStudio™ 5 Real-Time PCR System.

### Quantification and statistical analysis

Statistical analyses were performed using GraphPad Prism v.7.00 or Microsoft Excel softwares. Data are presented as mean and SEM. For all comparisons throughout our study, we performed Mann Whitney’s test or Student’s t-tests as specified on each figure legends. *p≤0.05, **p≤ 0.01, ***p≤0.001, ns = non-significant.

