## Supplemental figures for "Ingestion of *Bacillus cereus* spores dampens the immune response to favor bacterial persistence"

Hachfi et al.

### SUPPLEMENTAL FIGURES

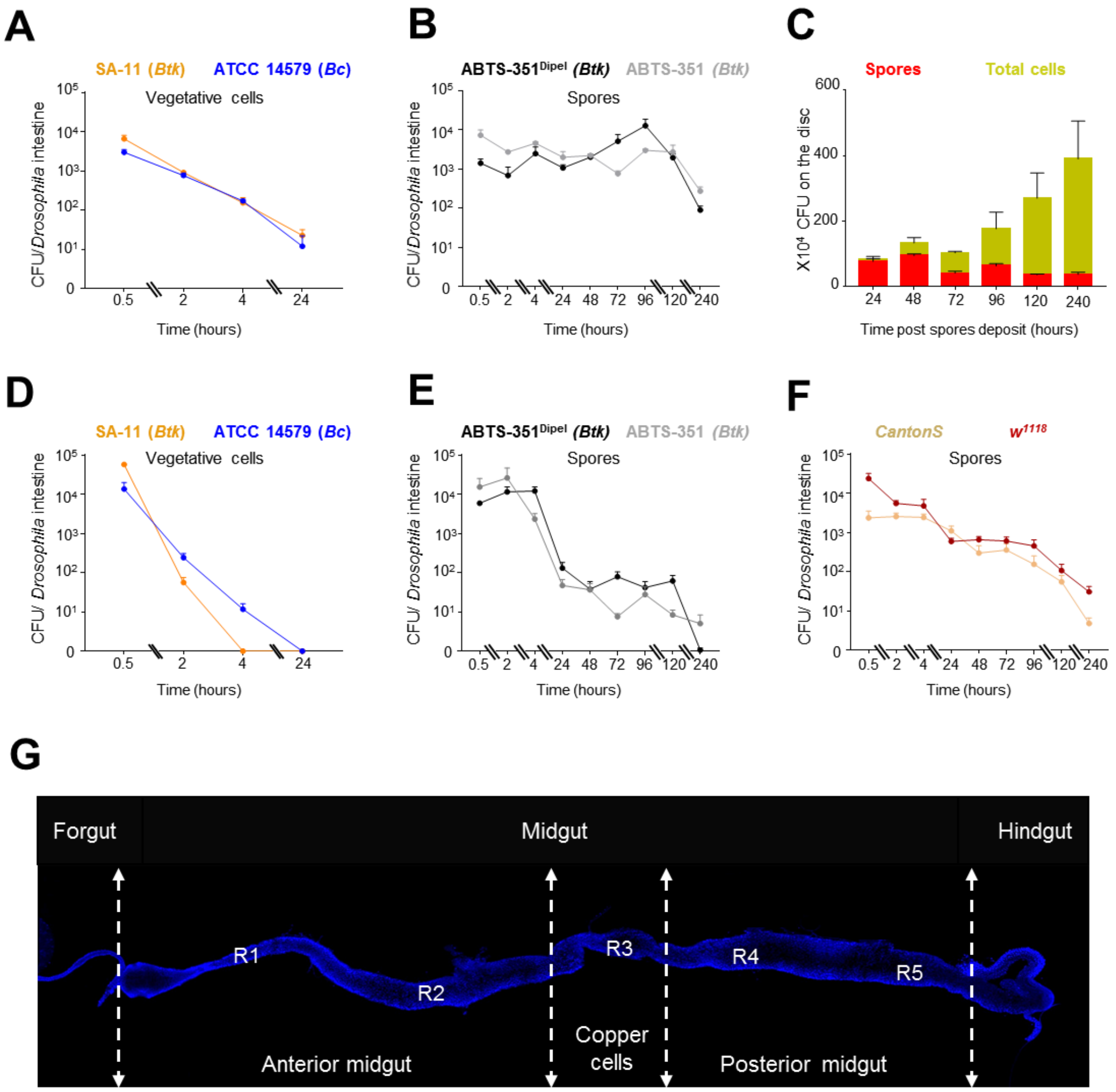


### Figure S1

**Figure S1: Spores of the *Bc* group persist in the *Drosophila* intestine**

**(A)** Vegetative cell numbers in Canton S flies' dissected midguts after continuous exposure. Flies were fed with SA-11 *(Btk)* vegetative cells (orange line) or ATCC 14579 (*Bc*) vegetative cells (blue line).

**(B)** Bacterial loads in dissected midguts after ABTS-351$^{\mathrm{Dipel}}$ (the commercial product, black line) and ABTS-351 ("homemade", gray line) spore continuous ingestion.

**(C)** Kinetic of the bacterial load on the feeding disc. The graph represents SA-11 (*Btk*) spore counts in red bars and SA-11 (*Btk*) total cell counts in green bars.

**(D)** Bacterial loads in dissected midguts of Canton S flies after acute exposure. Flies were fed with SA-11 (*Btk*, orange line) or ATCC 14579 (*Bc*, blue line) vegetative cells.

**(E)** Bacterial loads in dissected midguts after ABTS-351$^{Dipel}$ (the commercial product, black line) and ABTS-351 ("homemade", gray line) spore acute ingestion.

**(F)** Bacterial loads in dissected midguts from *Canton S* and $w^{1118}$ flies after SA-11 (*Btk*) spore acute exposure.

**(G)** *Drosophila melanogaster* midgut domains. Dotted lines represent where midguts have been sectioned in Figure 1C and Figure 2.

Error bars correspond to the SEM of at least three independent experiments.

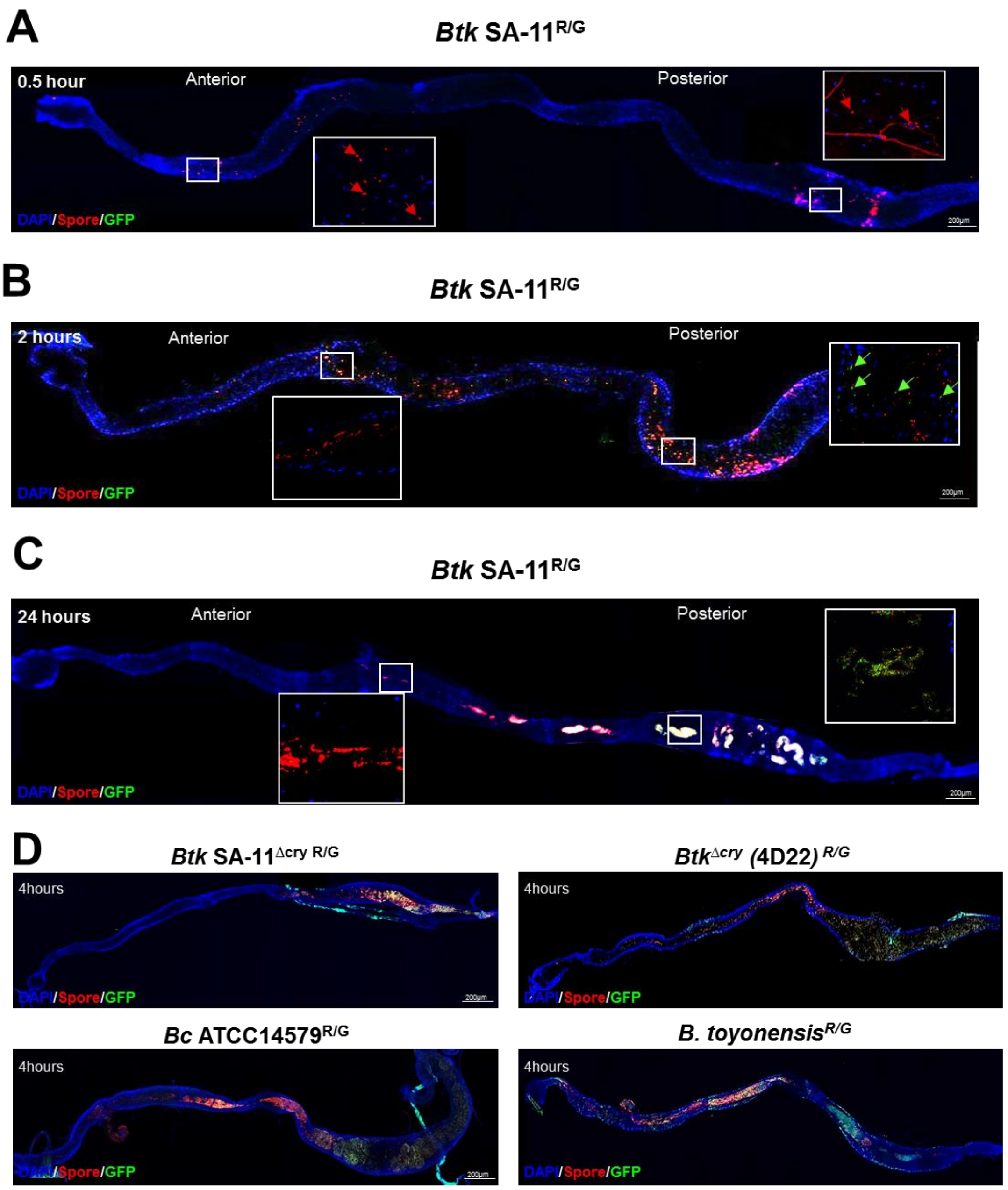


### Figure S2

**Figure S2: Spores of the *Bc* group germinate preferentially in the posterior midgut**

**(A-C**) Kinetic of SA-11$^{R/G}$ spore germination in the *Drosophila* intestinal midgut. Germinated vegetative cells (GFP) are observed from 4 hours onward post spore ingestion.

**(A)** Fluorescent image 0.5 hours post- SA-11$^{R/G}$ ingestion.

**(B)** Fluorescent image 2 hours post- SA-11$^{R/G}$ ingestion.

**(C)** Fluorescent image 24 hours post- SA-11$^{R/G}$ ingestion.

**(D)** *In vivo* spore$^{R/G}$ behavior from different strains of the *Bacillus cereus* group.

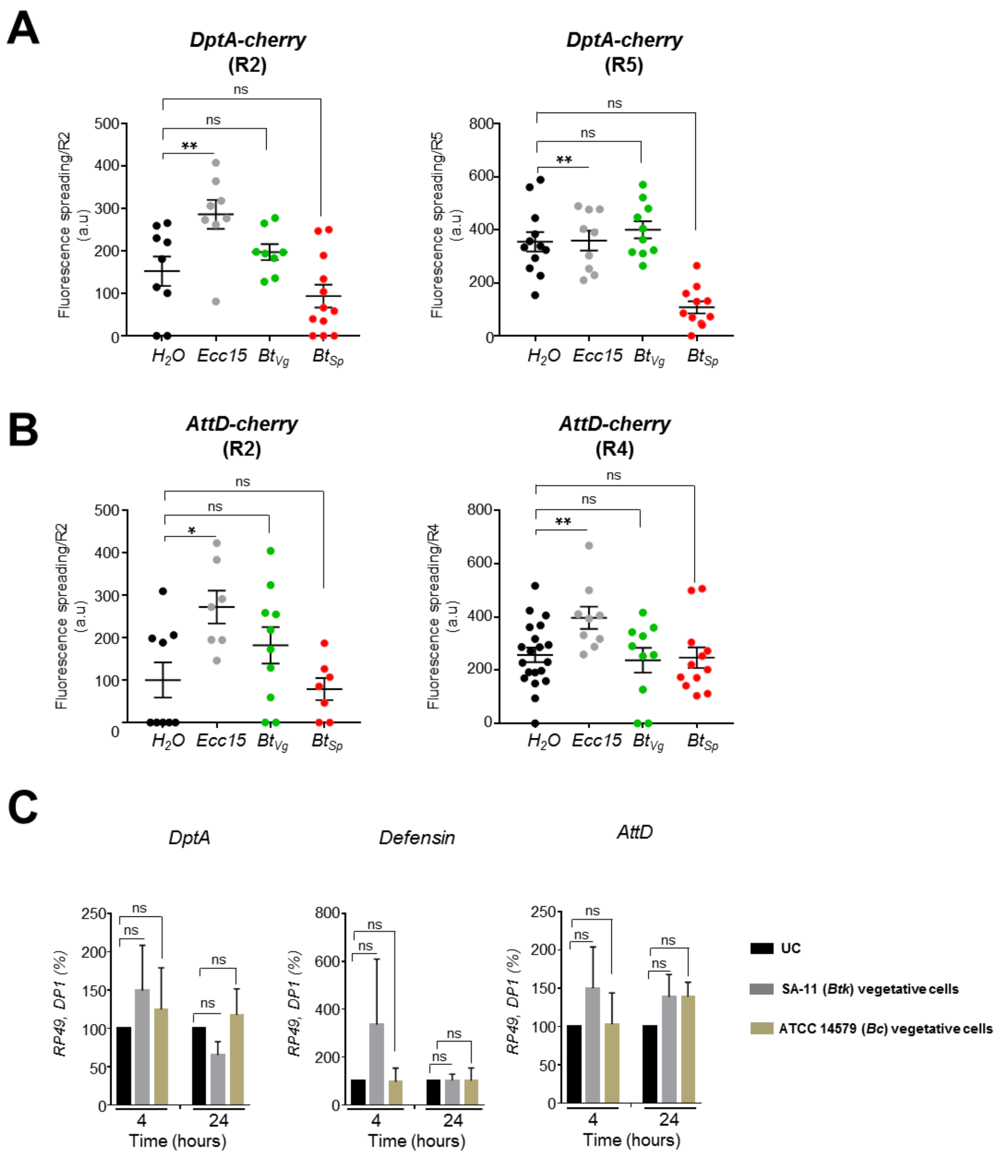


### Figure S3

**Figure S3: Spores do not trigger *Drosophila* midgut innate immune response**

**(A)** Spreading quantification of the *DptA-cherry* expression into the R2 (left panel) and R5 (right panel) regions.

**(B)** Spreading quantification of the *AttD-cherry* expression into the R2 (left panel) and R4 (right panel) regions.

**(C)** RT-qPCR analyses of *AMPs* expression 4 hours and 24 hours after acute feeding with *Bc* or SA-11 vegetative cells. UC corresponds to flies fed with water.

Mann-Whitney test was used to analyze data in A and B. Student's t-tests was used to analyze data in C. ns = non significant (p>0.05); *=p≤0.05; **= p≤0.01.

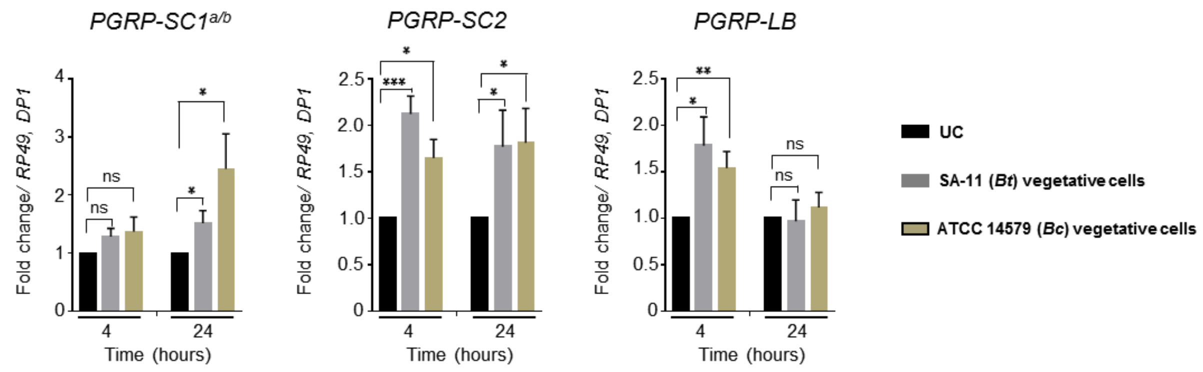


### Figure S4

**Figure S4: Amidases contribute to the intestinal persistence of spores**

RT-qPCR analysis of amidase expressions following *Bc* ATCC14579 or *Btk* SA-11 vegetative cells acute feeding.

UC corresponds to flies fed with water.

The results are shown as relative levels of expression. Data represent means ± SEM of at least three independent experiments. Student's t-tests was used: *=p≤0.05, * =p≤ 0.01, ***=p≤0.001, ns = non-significant (p>0.05).

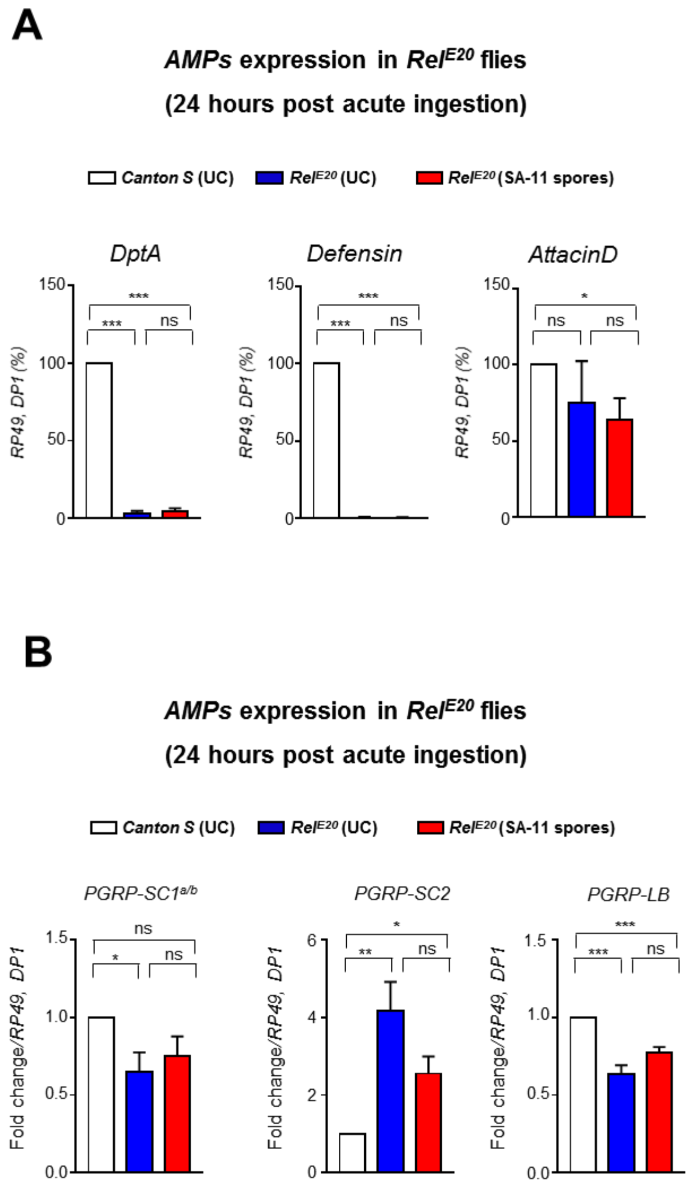


### Figure S5

**Figure S5: The Imd pathway contributes to the spore intestinal persistence**

**(A)** RT-qPCR analyses of *AMP* expressions in *Rel*$^{E20}$ mutant 24 hours after acute feeding with *Btk* SA-11 spores.

**(B)** RT-qPCR analyses of *amidase* expressions in *Rel*$^{E20}$ mutant 24 hours after acute feeding with *Btk* SA-11 spores.

UC corresponds to flies fed with water. The results are shown as relative levels of expression. Data represent means ± SEM of at least three independent experiments. Student's t-tests was used: *=p≤0.05, **=p≤ 0.01, ***=p≤0.001, ns = non-significant (p>0.05).

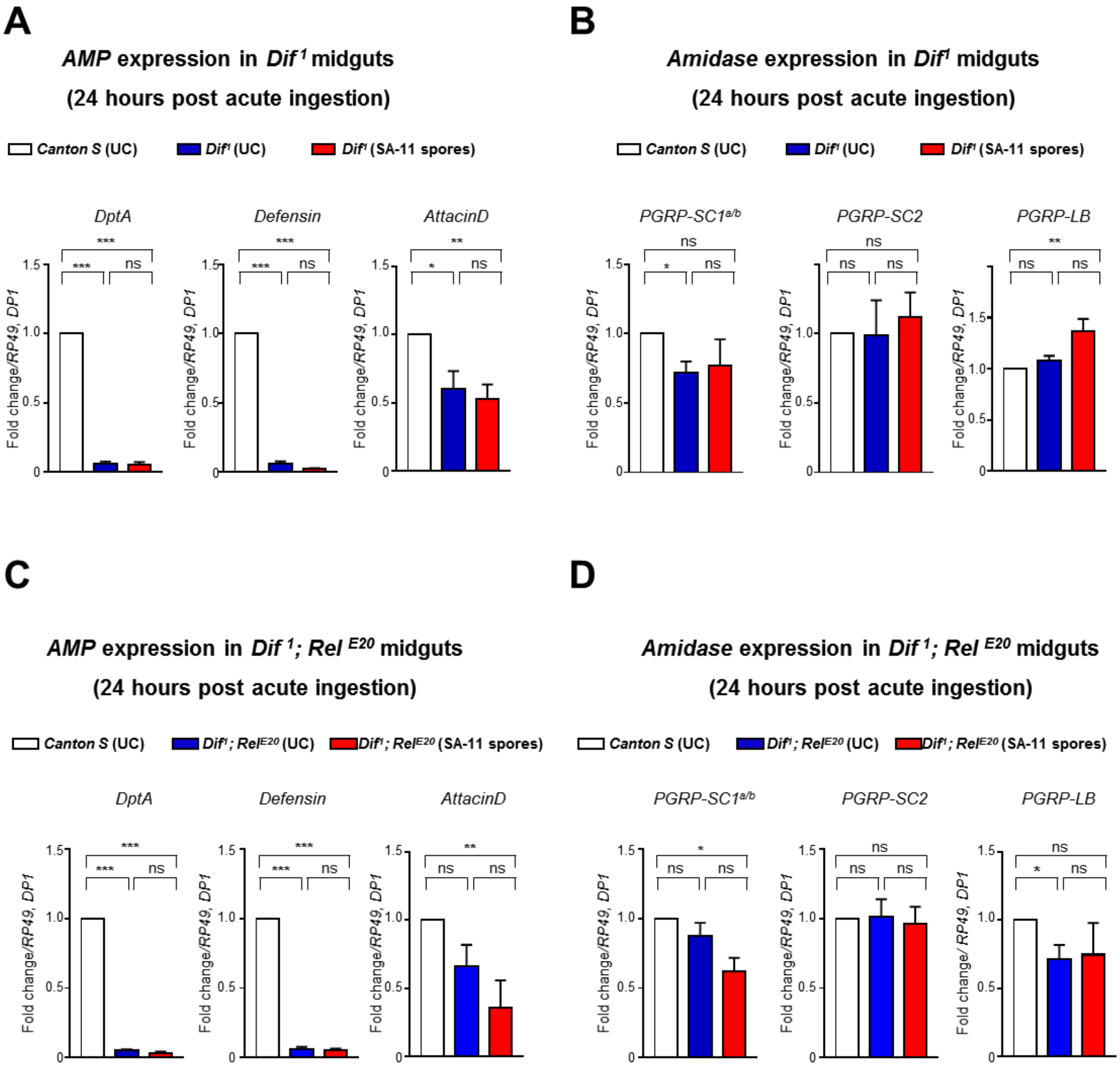


**Figure S6**

**Figure S6: Dif cooperates with Relish to modulate the intestinal immune response to spore ingestion**

**(A)** RT-qPCR analyses of *AMP* expressions in *Dif*$^1$ homozygous mutants 24 hours after acute feeding with *Btk* SA-11 spores.

**(B)** RT-qPCR analyses of *amidase* expressions in *Dif*$^1$ homozygous mutants 24 hours after

**(C)** RT-qPCR analyses of *AMP* expressions in *Dif*$^1$*; Rel*$^{20}$ homozygous double mutants 24 hours after acute feeding with *Btk* SA-11 spores.

**(D)** RT-qPCR analyses of *amidase* expressions in *Dif*$^1$*; Rel*$^{E20}$ homozygous double mutants 24 hours after acute feeding with *Btk* SA-11 spores.

UC corresponds to flies fed with water. The results are shown as relative levels of expression. Data represent means ± SEM of at least three independent experiments. Student's t-tests were applied: *=p≤0.05, **=p≤ 0.01, ***=p≤0.001, ns = non-significant (p>0.05).

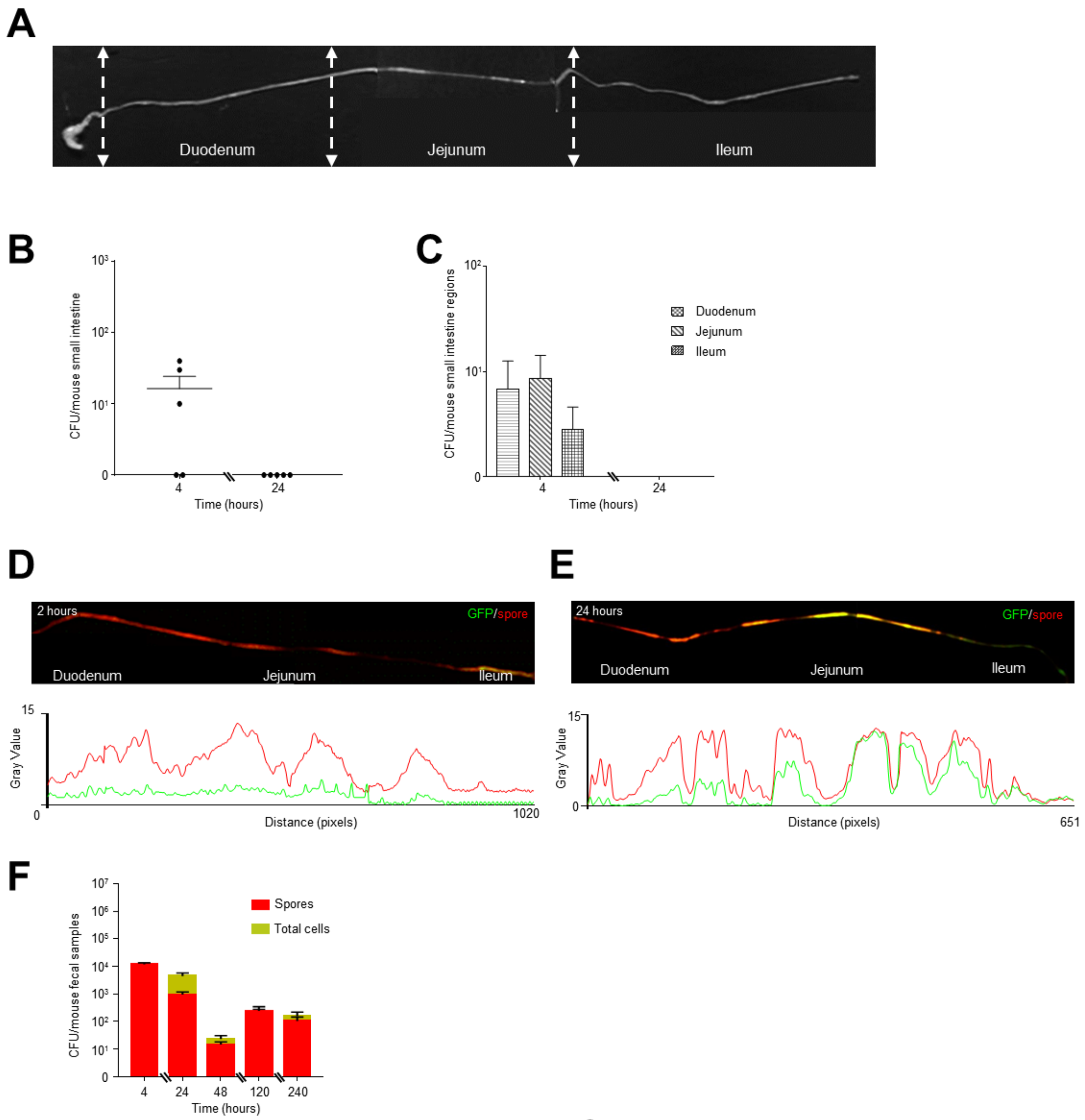


**Figure S7**

**Figure S7: *Btk* spores persist in the mouse intestine**

**(A)** Dissection scheme in mouse small intestines. Dotted lines represent where guts have been sectioned to isolate the different portions. Noteworthy, the zone delimited as the duodenum is larger than the reality. The duodenum is smaller but because of this absence of anatomical structure allowing the identification of the end of the duodenum, we have made the choice to split the small intestine in 3 equal parts.

**(B)** Bacterial loads in the small intestine after vegetative cells oral gavage.

**(C)** Localization of *Btk* SA-11 vegetative cells in the different part of the small intestines. Error bars correspond to the SEM of at least five independent experiments

**(D-E)** Monitoring of *Btk* SA-11$^{R/G}$ germination in small intestines 2 hours (A) and 24 hours (B) post gavage. Below are the plots of the average fluorescence intensity (represented as mean gray value) measured along mouse small intestines. Mice were continuously maintained in the same cage.

**(F)** Bacterial loads in mouse fecal samples after spore oral gavage. Mice were continuously maintained in the same cage.

**Movie1:** Time-lapse of SA-11$^{R/G}$ spores during germination.
